# The STROBE: a system for closed-looped optogenetic control of freely feeding flies

**DOI:** 10.1101/486050

**Authors:** Pierre-Yves Musso, Pierre Junca, Meghan Jelen, Damian Feldman-Kiss, Han Zhang, Rachel C.W. Chan, Michael D. Gordon

## Abstract

Manipulating feeding circuits in freely moving animals is challenging, in part because the timing of sensory inputs is affected by the animal’s behavior. To address this challenge in *Drosophila,* we developed the Sip-Triggered Optogenetic Behavior Enclosure (“STROBE”). The STROBE is a closed-looped system for real-time optogenetic activation of feeding flies, designed to evoke neural excitation coincident with food contact. We demonstrate that optogenetic stimulation of sweet sensory neurons in the STROBE drives attraction to tasteless food, while activation of bitter sensory neurons promotes avoidance. Moreover, feeding behavior in the STROBE is modified by the fly’s internal state, as well as the presence of chemical taste ligands. We also find that mushroom body dopaminergic neurons and their respective post-synaptic partners drive opposing feeding behaviors following activation. Together, these results establish the STROBE as a new tool for dissecting fly feeding circuits and suggest a role for mushroom body circuits in processing naïve taste responses.

## Introduction

*Drosophila melanogaster* has emerged as a leading model for understanding sensory processing related to food approach, avoidance, and consumption behaviors. However, although the gustatory system is recognized as mediating a critical final checkpoint in determining food suitability, much remains to be learned about the neural circuits that process taste information in the fly brain.

Like mammals, flies detect several taste modalities, each of which promotes food acceptance or rejection (Liman et al., 2014; Marella et al., 2006; Yarmolinsky et al., 2009). Taste compounds activate gustatory receptor neurons (GRNs) localized on the fly’s proboscis, legs, wings and ovipositor (Scott, 2018). Among the different classes of GRNs present, cells expressing the Gustatory Receptor (GR) Gr64f respond to sweet compounds and induce strong acceptance behavior. Conversely, GRNs labelled by Gr66a respond to bitter compounds and evoke avoidance (Dahanukar et al., 2001; 2007; Jiao et al., 2008; Kwon et al., 2014; 2011; Marella et al., 2006; Thorne et al., 2004; Wang et al., 2004). GRNs from the proboscis send direct axonal projections to the subesophageal zone (SEZ) of the fly brain (Ito et al., 2014; Rajashekhar and Singh, 1994; Scott, 2018; Stocker and Schorderet, 1981). Taste processing in the SEZ involves local modulatory interneurons (Chu et al., 2014; Pool et al., 2014), second-order neurons projecting locally or to other brain regions (Kain and Dahanukar, 2015; Kim et al., 2017; Yapici et al., 2016), motor neurons driving feeding subprograms (Gordon and Scott, 2009; Hampel et al., 2011; Manzo et al., 2012; Rajashekhar and Singh, 1994), and command neurons driving the complete feeding program (Flood et al., 2013).

Taste processing is not only involved in acute feeding events, but also in the formation of associative memories, which are aversive following exposure to bitter taste (Masek et al., 2015) or positive following sugar consumption (Tempel et al., 1983). Memory formation occurs mainly in a central brain structure called the Mushroom body (MB), composed of ~2000 Kenyon cells per hemisphere (Heisenberg et al., 1985). The MBs receive sensory information that is assigned a positive or negative output valence via coincident input from dopaminergic neurons (DANs) (Perisse et al., 2013; Waddell, 2010). Little is known about how taste information is relayed to the MBs, but Taste Projection neurons (TPNs) connected to bitter GRNs indirectly drive activation of the Paired Posterior Lateral cluster 1 (PPL1) DANs (Kim et al., 2017). PPL1 neurons signal punishment to MBs (Aso et al., 2012; 2010; Claridge-Chang et al., 2009) and are required for aversive taste memory formation (Kim et al., 2017; Kirkhart and Scott, 2015; Masek et al., 2015). Conversely, the Protocerebrum Anterior Medial cluster (PAM) cluster of DANs signals rewarding information and is involved in the formation of appetitive memories (Burke et al., 2012; Huetteroth et al., 2015; Liu et al., 2012; Yamagata et al., 2015). Although they have well-established roles in memory formation, PPL1 and PAM involvement in feeding has not been extensively investigated.

Kenyon cells and DANs make discrete connections with Mushroom body output neurons (MBONs), which project to protocerebral integration centers (Aso et al., 2014a; 2014b; Séjourné et al., 2011). MBONs are required for memory formation and retrieval (Aso et al., 2014b; 2014a; Bouzaiane et al., 2015; Felsenberg et al., 2017; Ichinose et al., 2015; Masek et al., 2015; Owald et al., 2015; Perisse et al., 2016; Plaçais et al., 2013; Séjourné et al., 2011; Takemura et al., 2017; Tanaka et al., 2008). In addition, MBONs can modulate innate behaviors such as taste sensitivity (Masek et al., 2015) or food seeking behavior (Tsao et al., 2018).

Manipulating neuron activity has become a powerful means of assessing neural circuit function. Silencing neuron populations in freely behaving flies is a straightforward way to determine their necessity in feeding, forcing the neurons in a chronic ‘off’ state to mimic a situation where the fly never encounters an activating stimulus (Fischler et al., 2007; Gordon and Scott, 2009; LeDue et al., 2015; 2016; Mann et al., 2013; Marella et al., 2012; Pool et al., 2014). However, gain-of-function experiments for feeding and taste, or any other actively sensed stimulus, are more complicated. Forcing a neuron into a stimulus- and behavior-independent “on” state can be difficult to interpret. The possible exception is activation of a neuron that elicits a stereotyped motor program, but even these situations are more easily interpreted in a harnessed fly where the effect of a single activation can be monitored (Chen and Dahanukar, 2017; Flood et al., 2013; Gordon and Scott, 2009; Marella et al., 2006; Masek et al., 2015). To effectively probe the sufficiency of neuron activation during feeding events, it would be ideal to temporally couple activation with feeding.

In a previous study, we harnessed the FlyPAD technology (Itskov et al., 2014) to develop a closed-loop system for real-time optogenetic activation of neurons during feeding behavior (Jaeger et al., 2018). This system, which we call the Sip-Triggered Optogenetic Behavior Enclosure (STROBE), triggers illumination of a red LED immediately upon detecting a fly’s interaction with one of two food sources in a FlyPAD arena. Here, we provide a more extensive characterization of the STROBE and its utility. Coincident activation of sweet GRNs with sipping on tasteless agar drives appetitive behavior, and bitter GRN activation elicits aversion. We show that these effects are modulated by starvation and can be inhibited by the presence of chemical taste ligands of the same modality. Activation of central feeding circuit neurons produces repetitive, uncontrolled sipping, demonstrating the ability of the STROBE to manipulate both peripheral and central neurons. We then demonstrate that activation of PPL1 neurons negatively impact feeding, while activating PAM neurons promotes it. Finally, in agreement with current olfactory memory models, activating DAN/MBON pairs from the same compartment drives opposing feeding behaviors.

## Results

### The STROBE triggers light activation temporally coupled with sipping

The FlyPAD produces a capacitance signal that increases when a fly physically interacts with the food on either of two sensors (“channels”) in a small arena (Figure 1a, Supplementary Figure 1a) (Itskov et al., 2014). This signal is then decoded *post hoc* by an algorithm designed to identify sipping events. The STROBE was designed to track the raw capacitance signal in real-time and trigger lighting within the arena during sips (Figure 1b). To achieve this, we built arena attachments that consist of a lighting PCB carrying two LEDs of desired colors positioned above the channels of the FlyPAD arenas. Each PCB is surrounded by a lightproof housing to isolate the arenas from other light sources (Supplementary Figure 1b-c).

**Figure 1.**
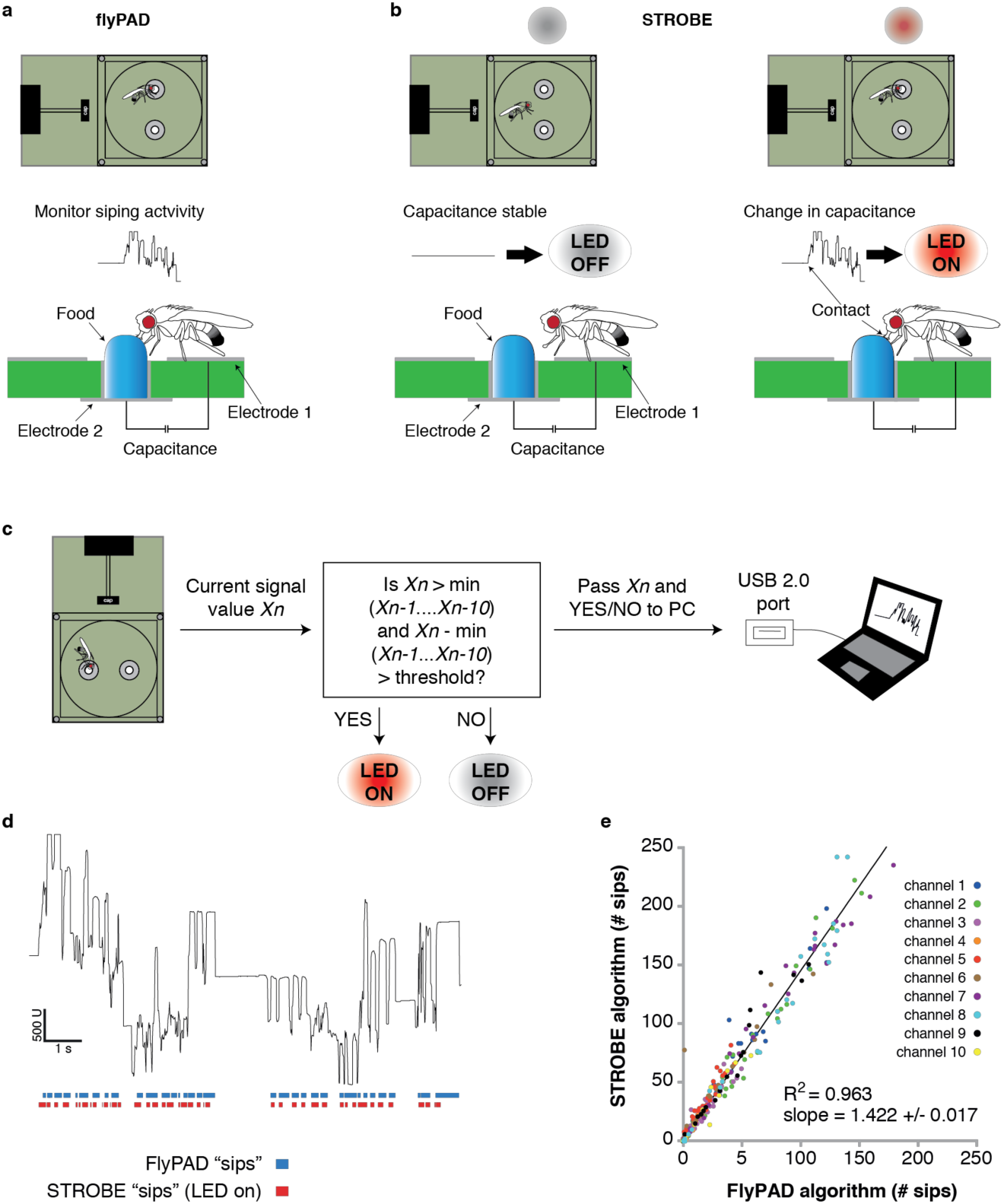
The STROBE setup. **a** Concept of the FlyPAD: The interaction between the fly’s proboscis and the food is detected as a change in capacitance between two electrodes: electrode 1, on which the fly stands, and electrode 2, on which the food is placed. **b** Concept of the STROBE: when the fly is not sipping, no change of capacitance is detected and the LED is OFF (left); when the fly sips, changes in capacitance turn the LED ON (right). **c** Flowchart of the STROBE signal processing algorithm. **d** Example of capacitance changes during a feeding bout, and the associated sips called by the FlyPAD (blue) and STROBE (red) algorithms. **e** Comparison of the sip numbers called by the FlyPAD and STROBE algorithms. Sips were counted in 1-minute bins across a 1-hour experiment for 10 different channels (5 arenas). Bins with no sips called by either algorithm were excluded from analysis.

In order to trigger optical stimulation with short latency upon sip initiation, we designed an algorithm that applies a running minima filter to the capacitance signal to detect when a fly is feeding. When a fly feeds, its contact with the capacitance plate generates a ‘step’, or rising edge in the capacitance signal. If this change surpasses a given threshold, then lighting is triggered. Because this algorithm is run on a field-programmable gate array (FPGA), the signal to lighting response transition times are on the order of tens of milliseconds, providing a nearly instantaneous response following the initiation of a sip. The system then records the state of the lighting activation system (on/off) and transmits this information through USB to the PC, where it is received and interpreted by a custom end-user program. This program displays the capacitance signals each fly arena in real-time, as well as its lighting state. It also counts the number of ‘sips’ (lighting changes) over the course of the experiment (Figure 1c; Supplementary Figure 1d-f).

To confirm that the STROBE algorithm detects sips in line with those detected by the original *post hoc* FlyPAD algorithm, we first used both algorithms to analyze the capacitance signal from a short (~11 s) feeding bout (Figure 1d). Visually, this showed us that each time a sip is detected with the FlyPAD algorithm, there is a sip detected at a similar time by the STROBE algorithm. However, we also noted that the STROBE algorithm called more sips than the FlyPAD algorithm. We confirmed these observations on a larger scale by examining the correlation between sip numbers detected by each algorithm in 1-minute bins across a full 1-hour experiment (Figure 1e). Here, we observed a strong correlation between the two (R^2^ = 0.963), with the STROBE algorithm detecting about 1.4 times the number of sips detected by the FlyPAD algorithm. This increased sip number is likely the consequence of the FlyPAD algorithm filtering out capacitance changes not adhering to certain criteria of shape and duration (Itskov et al., 2014). Since these parameters are, by definition, unknown at sip onset, the STROBE cannot use them as criteria. Thus, we expect a fraction of “sips” detected by the STROBE are actually more fleeting interactions with the food. Indeed, video of flies in the STROBE confirmed that leg touches also triggered light activation (Supplementary Movie 1). However, since flies detect tastes on multiple body parts, including the legs, these interactions are likely still relevant to taste processing and feeding initiation.

### Activation of GRNs modifies feeding behavior

To validate the utility of the STROBE, we first tested flies expressing CsChrimson, a red-light activated channel, in either sweet or bitter GRNs (Klapoetke et al., 2014). Flies were given the choice between two identical tasteless food options, one of which triggered light activation. Under these conditions, flies expressing functional CsChrimson in sweet neurons under control of *Gr64f-Gal4* showed a dramatic preference towards feeding on the light-triggering food (Figure 2a-e).

**Figure 2.**
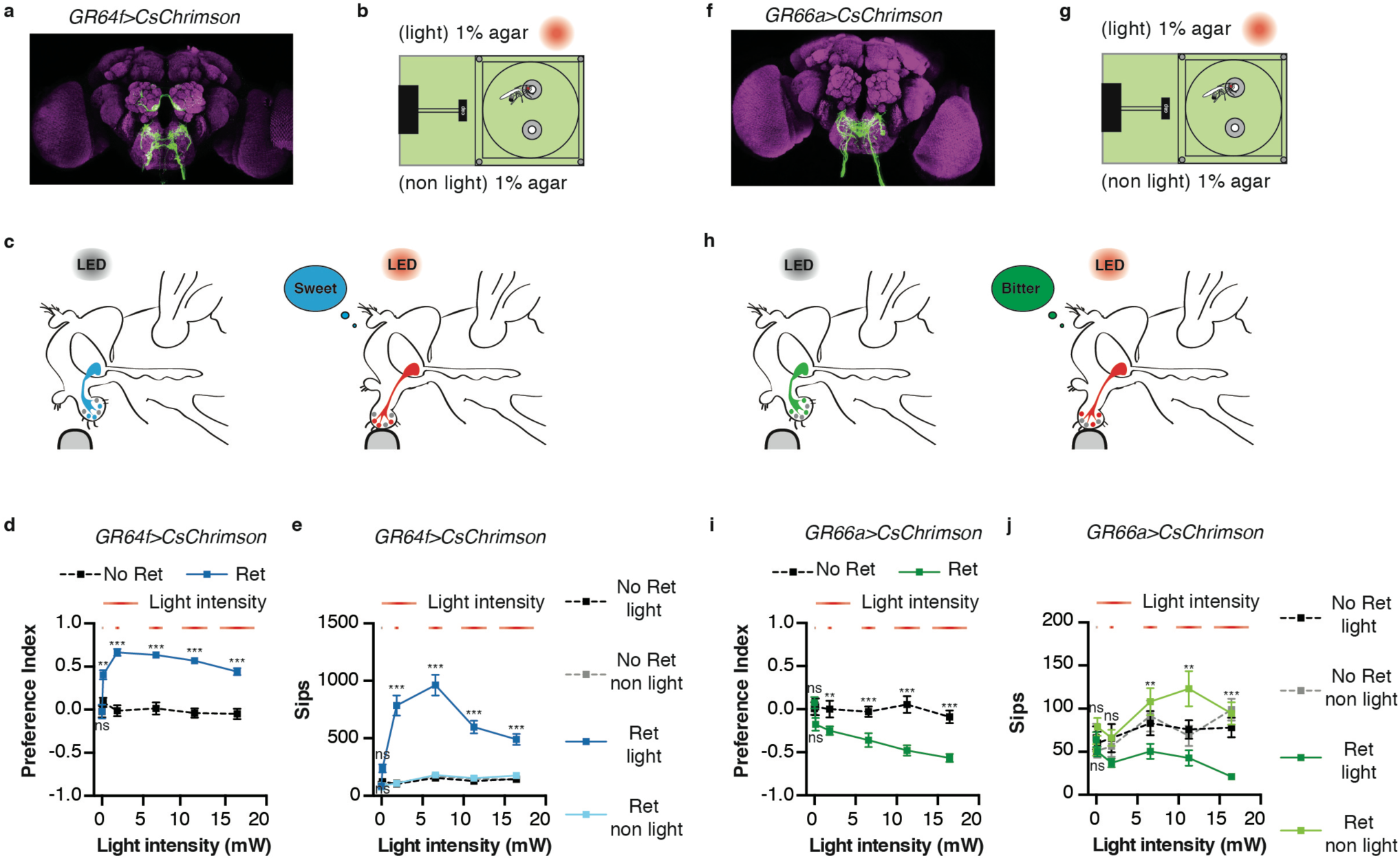
Activation of sweet and bitter sensory neurons drives feeding preferences in the STROBE. **a** Immunofluorescent detection of *UAS-CsChrimson.mVenus* driven by *Gr64f-GAL4*. **b** Experimental setup: both channels are filled with 1% agar, only one is associated to LED activation. **c** Schematic illustrating STROBE activation of sweet neurons. **d** Relationship between light intensity and preference for the light side of *Gr64f>CsChrimson* flies pre-fed retinal (blue squares; *n* = 32, 30, 37, 30, 34, 34) or not fed retinal (black squares; *n* = 32, 29, 38, 30, 38, 38). **e** Sip numbers for the experiment shown in c, demonstrating increased sips with increasing light intensity. **f** Expression of *UAS-CsChrimson.mVenus* driven by *Gr66a-GAL4*. **g** experimental setup: both channels contain plain 1% agar. **h** Schematic illustrating STROBE activation of bitter neurons. **i** Relationship between light intensity and preference for the light side of *Gr66a>CsChrimson* flies pre-fed retinal (green squares; *n* = 28, 23, 29, 26, 19, 26) or not fed retinal (black squares; *n* = 27, 17, 21, 24, 24, 29). **j** Sip numbers for the experiment in h. Values represent mean ± SEM. Statistical tests: *two-way ANOVA* and *Tukey post-hoc*; ns *p* > 0.05, ** *p* < 0.01, \*\*\**p* < 0.001.

Control flies of the same genotype that were not pre-fed all-*trans*-retinal, and thus carried non-functional CsChrimson, displayed no preference, demonstrating that the attraction to light is dependent on Gr64f neuron activation (Figure 2a-e). The increased sipping on the light-triggering side of the chamber was dependent on light intensity, with maximum sipping observed at 6.5 mW (Figure 2d).

As expected, flies expressing CsChrimson in bitter sensing neurons under the control of *Gr66a-Gal4* strongly avoided neuronal activation in the STROBE by sipping less on the light-triggering food source (Figure 2f-j). Once again, the behavioral response was intensity-dependent, with maximum suppression of sips occurring at the highest intensity tested (16.4 mW). Since the light intensity that gives the maximal effect for Gr64f and the Gr66a activation are 6.5 mW and 16.4 mW, respectively, we decided to use 11.2 mW as the light intensity for all further experiments.

### Behavioral impact of GRN activation is modulated by starvation

Starvation duration is well known to be a crucial parameter for feeding in flies – the longer flies are food deprived, the more they will accept sweet food and ingest it (Dus et al., 2013; 2011; Inagaki et al., 2012; 2014; Scheiner et al., 2004; Stafford et al., 2012). To determine if the STROBE could detect the physiological modulation of feeding behavior dependent on the internal state of the fly, we performed experiments in which starvation was manipulated (Figure 3a,b). In line with their behavior towards sugar, fed flies expressing CsChrimson in sweet neurons did not show a significant preference for the light-triggering food (Figure 3c). However, 12 hrs or more of starvation produced a clear preference toward the light side. This increased preference index is driven by a dramatic increase in sip number (Figure 3d). In contrast to its impact on sweet sensory neurons, starvation had no significant effect on the avoidance of light by bitter neuron activation (Figure 3e,f).

**Figure 3.**
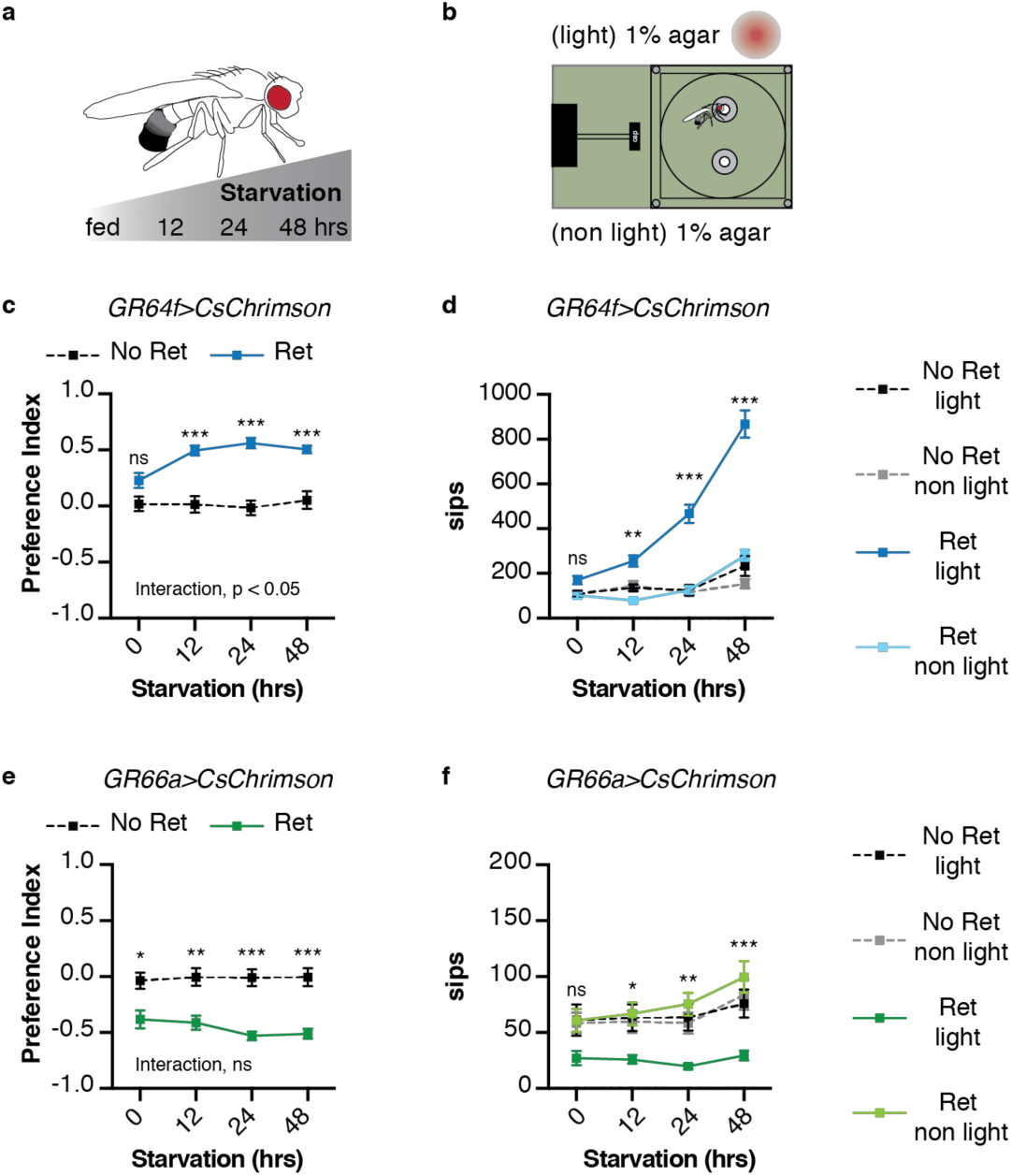
Behavioral impact of GRN activation is modulated by starvation. **a** Protocol: flies are subjected to increasing period of starvation (12 hrs, 24 hrs, 48 hrs) prior to the STROBE experiment. **b** Experimental setup: both channels contain plain 1% agar. **c-d** The effect of starvation on preference for the light side (c) and sip numbers (d) of flies expressing CsChrimson in sweet neurons (retinal flies: *n* = 30, 30, 25, 27; control flies: *n* = 29, 27, 21, 25). **e-f** The effect of starvation on preference for the light side (e) and sip numbers (f) of flies expressing CsChrimson in bitter neurons (retinal flies: *n* = 26, 23, 23, 24; control flies: *n* = 27, 27, 25, 25). Values represent mean ± SEM. Statistical tests: *two-way ANOVA* and *Tukey post-hoc*; ns *p* > 0.05, * p < 0.05, ** *p* < 0.01, \*\*\**p* < 0.001.

### Chemical taste ligands suppress the impact of light-induced attraction and avoidance

We next asked whether the presence of sweet or bitter ligands would interfere with light-driven behavior in the STROBE (Figure 4a,d; Supplementary Figure 2a,f). For example, if sugar is placed in both food options, will this reduce the salience of sweet GRN activation by light? Indeed, adding increasing concentrations of sucrose to both food options causes dose-dependent inhibition of the preference of *GR64f>CsChrimson* flies for the light-triggering food (Figure 4b). This change in preference is driven by progressively higher sipping on the non-light side, with relatively constant sip numbers on the light side as sucrose concentration increases (Supplementary Figure 2a-b). On the other hand, the addition of sucrose mildly enhanced the aversion driven by STROBE activation of Gr66a bitter neurons (Figure 4c; Supplementary Figure 2a,c). The same pattern is mirrored by the addition of the bitter compound denatonium to both sides: dose-dependent inhibition of the aversion shown by *Gr66a>CsChrimson* flies (Figure 4d-e; Supplementary Figure 2d-e), and little to no effect on the attraction of *Gr64f>CsChrimson* flies to the light side (Figure 4d,f; Supplementary Figure 2d,f).

**Figure 4.**
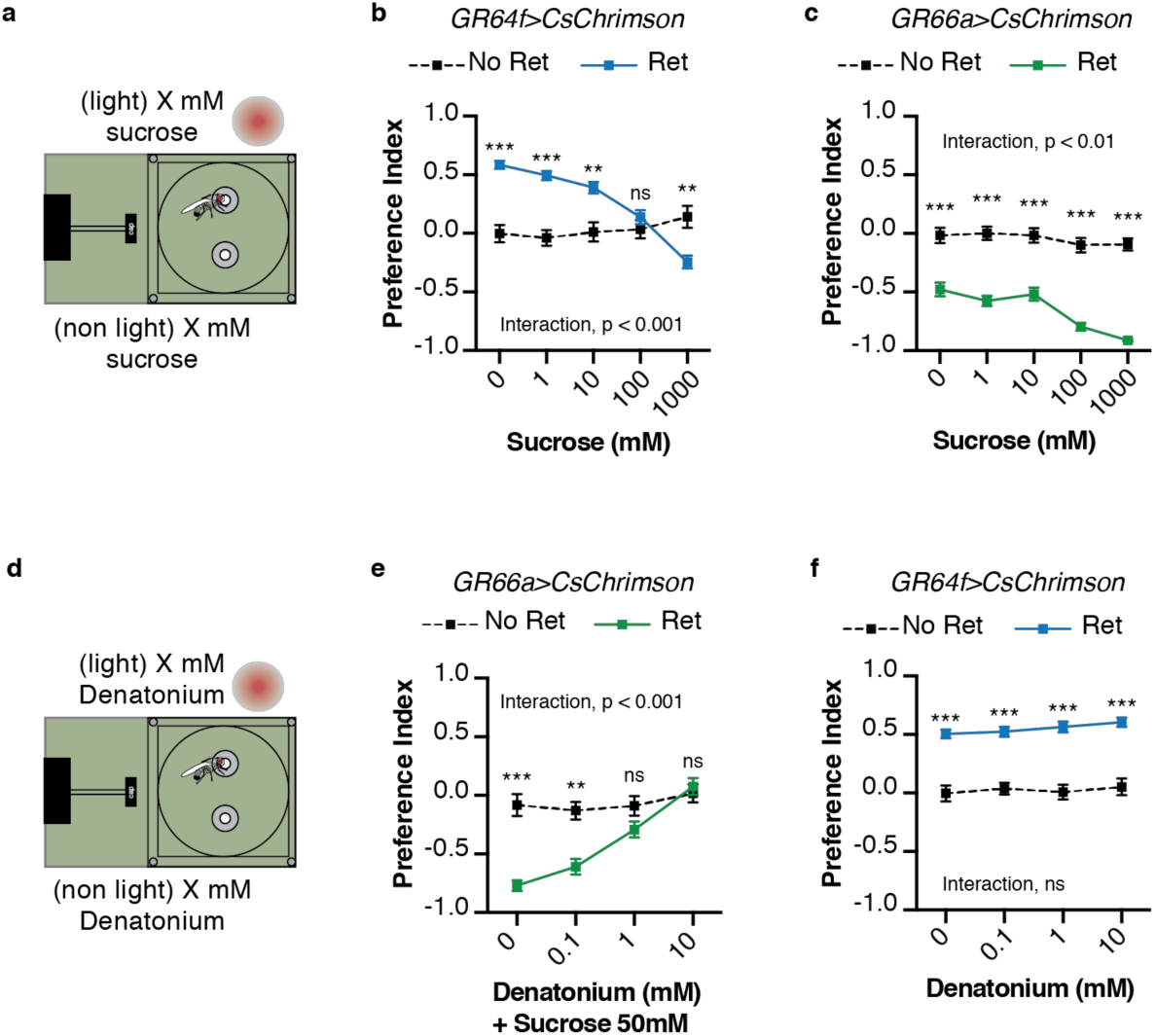
Chemical taste ligands suppress impact of light evoked GRN activity. **a** Experimental setup: both channels contain the same sucrose concentration (1, 10, 100, 1000 mM). **b-c** The effect of sucrose concentration on the preference of *Gr64f>CsChrimson* (b) or *Gr66a>CsChrimson* (c) for the light side (b, retinal flies: *n* = 28, 30, 30, 27, 30; control flies: *n* = 25, 26, 22, 25, 21; c, retinal flies: *n* = 28, 34, 34, 26, 46; control flies: *n* = 32, 40, 40, 37, 51). **d** Experimental setup: both channels contain the same denatonium concentration (0, 0.1, 1, 10 mM). For *Gr66a>CsChrimson* activation, both channels also contain 50 mM sucrose. **e-f** The effect of denatonium concentration on the preference of *Gr66a>CsChrimson* (e) or *Gr64f>CsChrimson* (f) for the light side (e, retinal flies: *n* = 25, 27, 30, 31; control flies: *n* = 26, 28, 27, 28; f, retinal flies: *n* = 30, 29, 29, 29; control flies: *n* = 30, 28, 27, 30). Values represent mean ± SEM; statistical tests: *two-way ANOVA*; *Tukey post-hoc*; ns: *p* > 0.05; \*\**p* < 0.01; \*\*\**p* < 0.001.

### Activation of the “feeding neuron” drives extreme sipping behavior

Can the STROBE could affect feeding behavior through the activation of central neurons, in addition to those in the periphery? Although the precise nature of higher-order taste circuits is still unclear, several neurons have been identified in the SEZ that influence feeding behavior (Chu et al., 2014; Inagaki et al., 2014; 2012; Jourjine et al., 2016; Kain and Dahanukar, 2015; LeDue et al., 2016; Marella et al., 2012; Pool et al., 2014; Yapici et al., 2016). One of them, the “feeding neuron” (Fdg), acts as a command neuron for the proboscis extension response, and shows activity in response to food stimulation only following starvation (Flood et al., 2013). Strikingly, *Fdg-GAL4>CsChrimson* flies placed in the STROBE with plain agar show an extremely high number of sips on the light side (Figure 5a-d; Supplementary Movie 2), resulting in a nearly complete preference for that side (Figure 5e-g), a preference retained when 100 mM sucrose is present (Figure 5h). Thus, the STROBE can effectively modulate feeding behavior via the activation of either peripheral or central neurons.

**Figure 5.**
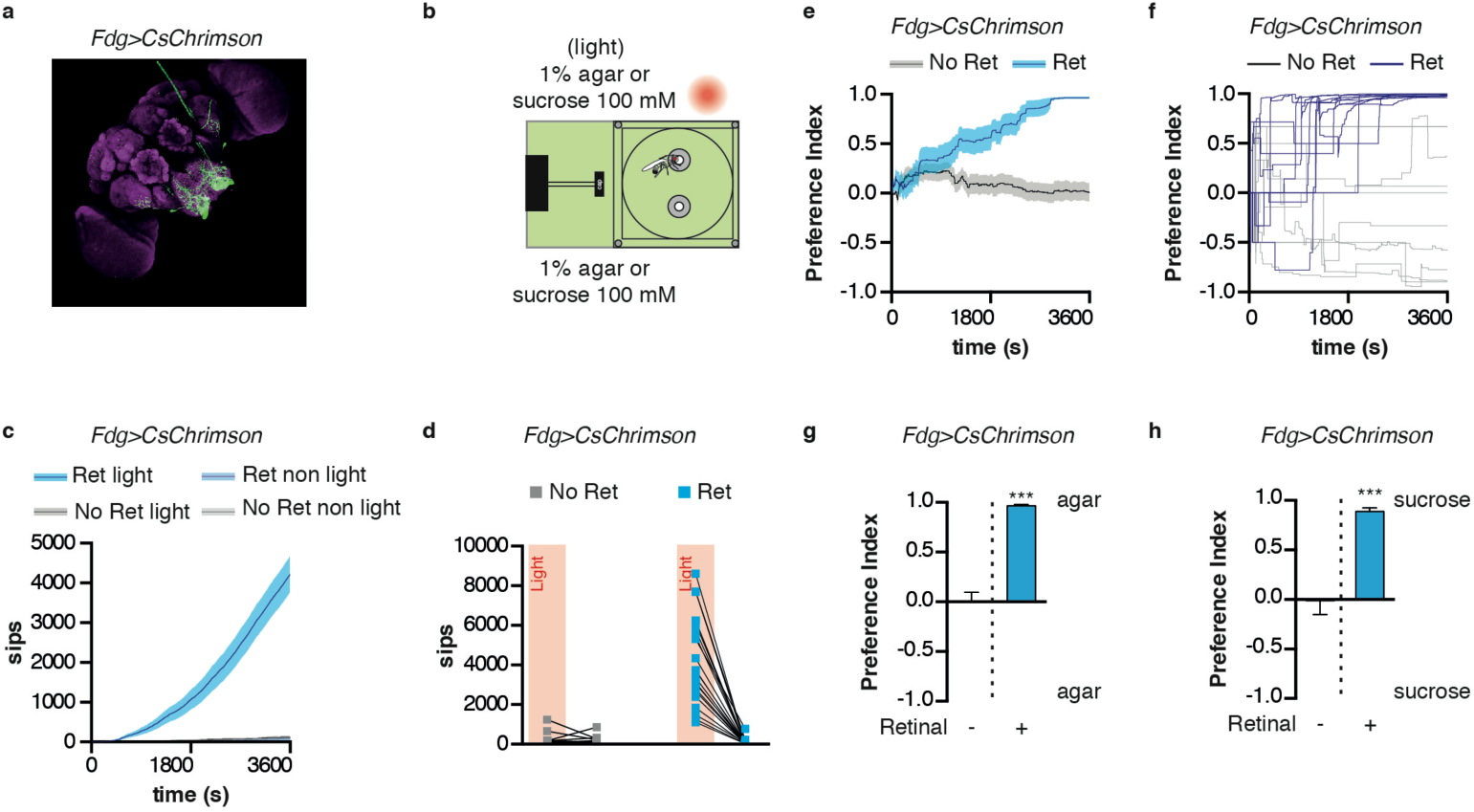
Activation of feeding command neurons elicits extreme sipping behavior. **a** Immunofluorescent detection of *Fdg>CsChrimson*. **b** Experimental setup: both channels are filled with 1% agar or 100 mM sucrose. **c-d** Cumulative sip numbers (c) for *Fdg>Chrimson* flies over the course of a 1-hour experiment (d) or for individual flies at the end of the experiment. **e-f** preference index for *Fdg>Chrimson* flies over the course of a 1-hour experiment averaged (e) or for ten individual flies (f). **g-h** Preference index for *Fdg>Chrimson* flies after one hour spent in the STROBE sipping on (g) agar or (h) sucrose 100 mM (g, retinal flies: n = 23; control flies: n = 33; h, retinal flies: n =18; control flies: n = 14). Values are means ± SEM. Statistical tests: *t*-test; \*\*\**p* < 0.001 in comparison between two groups.

### Manipulation of mushroom body extrinsic neurons modifies feeding behavior

The PAM and PPL1 clusters of dopaminergic extrinsic mushroom body neurons (DANs) are known to respond to taste inputs and mediate positive and negative reinforcement during learning (Burke et al., 2012; Das et al., 2014; Huetteroth et al., 2015; Kirkhart and Scott, 2015; Liu et al., 2012; Mao and Davis, 2009; Masek et al., 2015; Yamagata et al., 2015). Each DAN sends projections to a strikingly discrete compartment of the mushroom body, which contains the dendrites of specific Mushroom Body Output neurons (MBONs) (Aso et al., 2014b; 2014a). An emerging model is that the DAN/MBON pairs that innervate a specific MB compartment are opposite in valence, and the KC-MBON synapses in that compartment are depressed upon DAN activation (Aso et al., 2014b; Cohn et al., 2015; Felsenberg et al., 2017; Perisse et al., 2016; Séjourné et al., 2011; Takemura et al., 2017) (Figure 6a). We next asked whether activating DANs or MBONs coincident with sipping behavior would modulate feeding.

**Figure 6.**
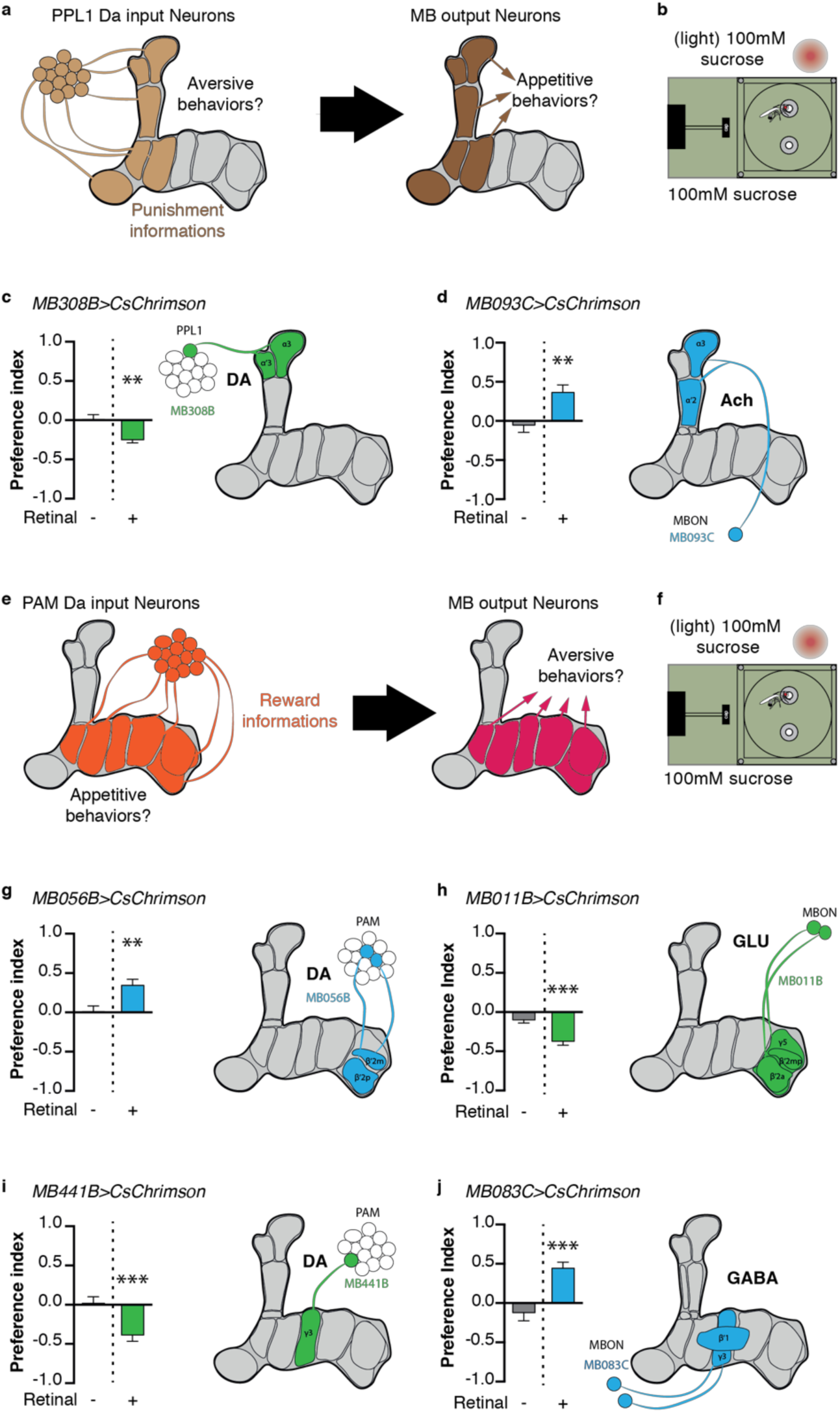
Manipulation of mushroom body extrinsic neurons modifies feeding behavior. **a** Model for PPL1 input to the MB and corresponding output: PPL1 signals punishment and should drive aversive behavior, while corresponding MBONs should be appetitive. **b** Experimental setup: both channels contain 100 mM sucrose in 1% agar. **c** Preference of flies expressing CsChrimson in PPL1 neurons α3,α’3 (*MB308-GAL4*) (retinal flies: *n* = 29; control flies: *n* = 27). **d** Preference of flies expressing CsChrimson in MBON α3,α’2 neurons post-synaptic to PPL1 neurons (*MB093C-GAL4*) (retinal flies: *n* = 26; control flies: *n* = 18). **e** Model for PAM input to the MB and corresponding output: PAM signals reward information and should drive appetitive behavior while corresponding MBON should be aversive. **f** Experimental setup: both channels contain 100 mM sucrose in 1% agar. **g** Preference of flies expressing CsChrimson in PAM β2,β’2 neurons (*MB056B-GAL4*) (retinal flies: *n* = 24; control flies: *n* = 24). **h** Preference of flies expressing CsChrimson in MBON neurons post-synaptic to PAM β2,β’2 neurons (*MB011B-GAL4*) (retinal flies: *n* = 25; control flies: *n* = 27). **i** Preference of flies expressing CsChrimson in PAM γ3 neurons (*MB441-GAL4*) (retinal flies: *n* = 25; control flies: *n* = 24). **j** Preference of flies expressing CsChrimson in MBON neurons post-synaptic to PAM γ3 neurons (*MB083C-GAL4*) (retinal flies: *n* = 25; control flies: *n* = 25). Values are means ± SEM. Statistical tests: *t*-test; \*\**p* < 0.01; \*\*\**p* < 0.001 in comparison between two groups.

Since PPL1 neurons signal punishment or aversive information to the MBs (Figure 6a; (Aso et al., 2012; 2010; Das et al., 2014; Kirkhart and Scott, 2015; Masek et al., 2015)), their activation in the STROBE is predicted to drive avoidance of the light-triggering food (Figure 6a). Flies expressing CsChrimson in the α3/α’3 subset of PPL1 dopaminergic neurons (*MB308B-GAL4*) show a negative preference towards light when 100 mM sucrose is present in both options, but not if the food is tasteless agar alone (Figure 6b-c; Supplementary Figure 3a-c). Interestingly, performing the same experiment with flies expressing CsChrimson in a broader subset of PPL1 neurons (*MB065B-GAL4*) leads to stronger avoidance in the presence of 100 mM sucrose as well as plain agar (Supplementary Figure 3a-c). Interestingly, activation of MBONs (*MB093C-GAL4* and *MB026B-GAL4*) post-synaptic to PPL1 produces strong attraction in the presence of 100 mM of sucrose (Figure 6d; Supplementary Figure 3a-c) or plain agar (Supplementary Figure 3a-c).

PAM neurons signal appetitive reward to the MBs (Burke et al., 2012; Huetteroth et al., 2015; Lin et al., 2014; Liu et al., 2012; Yamagata et al., 2015). Following the same principle, PAM activation should drive appetitive behavior, while stimulation of MBONs within the same compartment is predicted to elicit aversion (Figure 6e). Indeed, expressing CsChrimson in PAM β2m,β’2p neurons (*MB056B-GAL4* and *MB301B-GAL4*) leads to attraction in the presence of sucrose (Figure 6f,g; Supplementary Figure 4a-c) or plain agar (Supplementary Figure 4a-c). On the other hand, activating the corresponding β2m,β’2p,γ5 MBONs (*MB011B-GAL4* and *MB210B-GAL4*) produces avoidance (Figure 6h, Supplementary Figure 4a-c).

Another subset of PAM neurons, targeting the γ3 compartment, has recently been shown to be post-synaptic to neuropeptidergic Allatostatin-A neurons, which signal satiety state through inhibition (Hergarden et al., 2012; Yamagata et al., 2016). Interestingly, activation of PAM g3 neurons (*MB441B-GAL4* and *MB195B-GAL4*) in the STROBE is aversive (Figure 6i; Supplementary Figure 5a-c), while activation of the corresponding β’1,γ3 MBONs (*MB083C-GAL4* and *MB110C-GAL4*) is attractive (Figure 6j; Supplementary Figure 4a-c). Thus, PAM neurons targeting different MB compartments can be either attractive or aversive in the context of feeding.

## Discussion

Leveraging real-time data from the FlyPAD, we built the STROBE to tightly couple LED lighting with sipping events, thereby allowing us to optogenetically activate specific neurons during feeding. To demonstrate the STROBE’s utility, we showed that flies expressing CsChrimson in sweet or bitter GRNs display attraction and aversion, respectively, toward food that triggers LED lighting. Activation of central feeding command neurons also produces a dramatic enhancement of feeding behavior. Finally, we probed the effects of manipulating mushroom body input and output neurons, and demonstrated that activating DANs and MBONs within the same MB compartment generally produced opposing effects on feeding.

The primary advantage of the STROBE over existing systems for neural activation during fly feeding is its temporal resolution, which provides two important benefits. First, activating neurons while the fly is choosing to interact with one of two available food sources allows us to explore the impact of neural activation on food selection in a way that is impossible with chronic activation mediated by either temperature or light. Second, by tightly coupling activation with food interaction events, light-driven activity from the STROBE should more closely mimic the temporal dynamics of taste input. Conceptually, these advantages are similar to those achieved by expression of the mammalian TRPV1 in taste sensory neurons and lacing food with capsaicin (Caterina et al., 1997; Chen and Dahanukar, 2017; Marella et al., 2006). Importantly, however, the STROBE allows activation of either peripheral or central neurons.

Timing of activation also distinguishes the STROBE from another recently described optogenetic FlyPAD (Steck et al., 2018). The implementation of sip detection and light triggering on the FPGA allows the STROBE to trigger LED activation with minimal latency. Thus, neural activation is tightly locked to sip onset and offset, allowing the manipulation of circuits during active food detection. In contrast, the system described by Steck and colleagues (2018) carries out sip detection and light control on the USB-connected computer. The benefits of this strategy are the ability to implement a more complex feeding detection algorithm, and more flexible control of the lighting response timing. However, the tradeoff is longer and more variable latencies of LED activation. Each of these systems may have specific advantages, depending on the application. While they have not been directly compared, it is likely that tight temporal coupling of the STROBE to sips will be more useful for studying the effects of acutely activating core taste and feeding circuit neurons, while the longer, adjustable, light pulse from the optogenetic FlyPAD may be better for silencing neurons or activating reinforcement circuits.

Interestingly, a similar closed-loop optogenetic setup has recently been developed for mice. Lick-triggered blue light stimulation of the tongue enhanced licking in water-deprived mice expressing Channelrhodopsin2 in acid sensing taste receptor cells (Zocchi et al., 2017). Thus, the same principle is able to reveal important insight into consummatory behaviors in multiple animals.

Although optogenetic neuronal activation is artificial, light-driven behavior in the STROBE shows some important properties that mimic natural feeding. Starvation is known to directly increase feeding behavior on sweet food in flies (Dus et al., 2013; 2011; Inagaki et al., 2014; 2012; Scheiner et al., 2004; Stafford et al., 2012). As expected, increasing starvation led to an increased sipping on the light side for flies expressing CsChrimson in sweet sensory neurons, demonstrating behavior related to artificial activation of sweet sensory neurons is regulated by the flies’ internal state.

We also showed that the behavioral impact of sweet and bitter GRN activation in the STROBE could be abolished by the presence of natural taste ligands. Interestingly, this property did not hold true for attraction mediated by PAM or appetitive MBON activation, which was similar in the presence or absence of sugar. This may suggest that sweet taste input and PAM or MBON activation drive attraction via parallel circuits, producing an additive effect when both are present. It is also notable that that flies preferred 1 M sucrose alone over 1M sucrose coupled to optogenetic activation of sweet GRNs (Figure 4b). We suspect that optogenetic activation of sweet GRNs in the STROBE plateaus below the excitation achieved with 1 M sucrose, and somehow prevents further activation by very high sugar concentrations.

One interesting question is whether the valence observed from GRN activation in the STROBE is mediated by hedonics or effects on the feeding program itself. For example, sweet neuron activation is thought to carry appetitive hedonics, and therefore the flies may continue feeding because consequent light activation of Gr64f neurons is somehow pleasurable. On the other hand, these neurons also initiate feeding (and conversely, activation of Gr66a neurons terminates it). Thus, it is possible that each sip evokes light-driven activation of a subsequent sip, and so on, creating a positive feedback loop. This is undoubtedly true of Fdg neuron activation, which is known to initiate a complete feeding sequence, likely downstream of any hedonic effects (Flood et al., 2013). Flies appear to become “trapped” in a feeding loop until the end of the experiment, suggested by the very high number of sips evoked (see Figure 5).

We observed that activation of high order neurons such as Mushroom body DANs and MBONs also modulate sipping events in the STROBE. PPL1 DANs project mainly onto the vertical lobes of MBs have been described as signaling punishment information to the MBs for memory formation (Aso et al., 2012; 2010; Das et al., 2014; Kirkhart and Scott, 2015; Masek et al., 2015). In the present study, we observe that paired activation of PPL1 with food contact leads to an acute avoidance of the food. On the other hand, PAM DANs that signal reward information to the MBs (Burke et al., 2012; Huetteroth et al., 2015; Lin et al., 2014; Liu et al., 2012; Yamagata et al., 2015) lead to increased interactions with the light-triggering food source. Moreover, MBON activation from the same compartment drives opposing feeding behavior when compared to corresponding DAN stimulation (see Figure 6; Supplementary Figures 3-5). This relationship supports the current model that DAN activity depresses KC to MBON synapses in their respective compartment (Cohn et al., 2015; Felsenberg et al., 2017; Perisse et al., 2016; Séjourné et al., 2011; Takemura et al., 2017). It is also interesting to note that the PAM neurons are not universally positive. PAM γ3 neurons display activity in response to electric shocks (Cohn et al., 2015) and are silenced upon sucrose stimulation (Cohn et al., 2015; Yamagata et al., 2016). Our findings that PAM γ3 activation drives aversive feeding behavior supports these neurons conveying a negative valence onto the MB (see Figure 6, Supplementary Figure 5).

Although the role of DANs in feeding behavior remains unclear, accumulating evidence suggests that MBONs can modulate innate behavior such as taste sensitivity (Masek et al., 2015), naïve response to odors (Owald et al., 2015), place preference (Aso et al., 2014b) or food seeking behavior (Tsao et al., 2018). Could the effect of DANs in the STROBE be mediated by a learning-driven process? This seems unlikely given our current experimental design as both food options were always identical, and thus there would be no cues to associate with appetitive or aversive DAN stimulation. We think it is more likely that the same reward or punishment signals that underlie memory formation also acutely modify feeding behavior. However, the possibility of pairing circuit activation with specific food cues may offer a new paradigm for studying food memories, and neuronal activation by self-administration opens a potential new avenue for the study of addiction.

## Methods

### STROBE System

The STROBE system consists of a field programmable gate array (FPGA) controller attached to a multiplexor board, adaptor boards, fly arenas equipped with capacitive sensors and lighting circuits. The hardware, with the exception of the lighting circuit units, is based on the FlyPAD design (Itskov et al., 2014). Each fly arena is paired with a lighting circuit and an opaque curtain (to prevent interference from external light). This pair will be referred to as a fly chamber unit. The entire system accommodates 16 fly chamber units (16 fly arenas and 16 lighting circuits), through 8 adaptor boards. The FPGA used is a Terasic DEV0-Nano mounted onto a custom-made multiplexor board.

The multiplexor board is one of the intermediate connection components between the fly chambers and the FPGA controller. The multiplexor board has eight 10-pin ports each of which facilitate communications between two fly chambers and the FPGA controller. The board also has a FTDI module allowing data transfer over serial communications with a computer. The other intermediate connection component is the adaptor board which connects on one side to the multiplexor board via a 10-pin line, and splits the 10-pin line from the multiplexor board into four 10-pin ports which connects to two fly arenas and two lighting circuits. The fly arena consists of two annulus shaped capacitive sensors and a CAPDAC chip (AD7150BRMZ) that the main multiplexer board communicates with to initiate and collect data (and ultimately to stop collecting data). The CAPDAC interprets and converts capacitance data from the two sensors on the fly-arena to a digital signal for the FPGA to process (Itskov et al., 2014).

The lighting circuit consists of a two-pin connector to receive power from an external power supply, a 10-pin connector to receive signals from the FPGA controller via the intermediate components, a 617 nm light emitting diode (LUXEON Rebel LED – 127lm @ 700mA; Luxeon Star LEDs #LXM2-PH01-0060), two power resistors (TE Connectivity Passive Product SMW24R7JT) for LED current protection, and two metal oxide semiconductor field effect transistors (MOSFETs; from Infineon Technologies, Neubiberg, Germany, IRLML0060TRPBF) allowing for voltage signal switching of the LEDs.

When a fly performs a sip and triggers a high signal on a capacitive sensor, the CAPDAC on the fly arena propagates a signal via the multiplexor to the FPGA controller. The FPGA processes the capacitive sensor signal, decides a legitimate sip was detected and sends a high signal through the multiplexor to the MOSFET of the lighting circuit. The MOSFET then switches its lighting circuit on, allowing current to flow and turning on the monocolor LED positioned directly above the capacitive sensor. The process for determining a legitimate sip is described next.

In order to trigger optical stimulation with short latency upon sip initiation, we designed a running minima filter that operates in real-time to detect when a fly is feeding. We implemented this filter by modifying the state machine on the FPGA. When a fly feeds, its contact with the capacitance plate generates a ‘step’, or rising edge in the capacitance signal. Our filter determines the minimum signal value in the last 100 ms and checks whether the current signal value is greater than this minimum. It further determines whether the current value is sufficiently large to be considered a rising edge relative to this minimum, based on a threshold set to exceed noise. If both these conditions are true, then the filter will prompt the lighting activation system to activate the LED (or keep it on if it is already on).

By design, this means that the control system will send a signal to deactivate the lighting upon the falling edge of the capacitance signal, or if the capacitance signal has plateaued for 100ms, whichever comes sooner. At this point, a low signal is sent to the MOSFET which pinches off the current flowing through the lighting circuit, turning off the light. The signal to lighting response transition times are on the order of tens of milliseconds, providing a nearly instantaneous response.

After each lighting decision (on/off/no change), the system will then automatically record the state of the lighting activation system (on/off) and transmit this information through USB to the computer, where it is received and interpreted by a custom end-user program (built using Qt framework in C++) which can display and record both the activation state and signal measured by the STROBE system for each channel of every fly arena, in real-time.

All STROBE design materials are available as a supplemental download.

All STROBE software is available for download from Github:

FPGA code: https://github.com/rcwchan/STROBE-fpga

All other code: https://github.com/rcwchan/ STROBE_software/

### Fly strains

Fly stocks were raised on standard food at 25 °C and 60% relative humidity under a 12:12 h light:dark cycle. For neuronal activation experiments we used the *20XUAS-IVS-CsChrimson.mVenus* (in attP40 insertion site) from the Bloomington *Drosophila* Stock Center (stock number: 55135). Specific GRN expression was driven using *Gr64f-GAL4* (Dahanukar et al., 2007) and *Gr66a-GAL4* (Wang et al., 2004). *GMR81E10*-*Gal4* was used for expression in Fdg neurons (Jenett et al., 2012; Pool et al., 2014). All MB split-GAL4 lines (*MB011B-GAL4*; *MB026B-GAL4*; *MB056B-GAL4*; *MB065B-GAL4*; *MB083C-GAL4*; *MB093C-GAL4*; *MB10C-GAL4*; *MB195B-GAL4*; *MB210B-GAL4*; *MB308B-GAL4*; *MB310B-GAL4*; *MB441B-GAL4*) were described in a previous study (Aso et al., 2014b) and obtained directly from Janelia. The expression patterns of the lines from the Flylight collections are available from the Flylight project websites.

### Fly preparation and STROBE experiments

After eclosion, adult female flies were kept for several days in fresh vials containing standard medium, and were then transferred at 25 °C into vials covered with aluminum foil containing 1ml standard medium (control flies) or into vials containing 1ml standard medium mixed with 1mM of all-*trans*-retinal (retinal flies) for 2 days. Then flies were transferred to vials covered with aluminum foil containing 1ml of 1% agar (control flies) or into vials containing 1ml of 1% agar mixed with 1mM of all-*trans*-retinal (retinal flies) for 24 hours.

For the starvation curve experiment (Figure 3), flies were transferred into vials containing 1 ml of standard medium +/-all-*trans*-retinal for 24 hours (fed group); or 1ml of 1% agar +/-all trans-Retinal for 12-24-48 hours.

All flies were 5-9 days old at the time of the assay. Both channels of STROBE chambers were loaded with 4 μl of 1% agar with or without sucrose (0, 1, 10, 100, 1000 mM) or denatonium (0, 0.1, 1, 10 mM). For aversive assays using denatonium, 50 mM sucrose was also added to increase basal sips number.

Acquisition on the STROBE software was started and then single flies were transferred into each arena by mouth aspiration. Experiments were run for 60 minutes, and the preference index for each fly was calculated as: (sips from Food 1 – sips from Food 2)/(sips from Food 1 + sips from Food 2). The red LED is always associated to the left side (Food 1). For temporal curves, data are pooled within 1s time-period.

Sucrose, denatonium, agar and all-*trans*-retinal were obtained from Sigma-Aldrich.

For experiments done in figure 2, light intensity used are 0, 0.12, 1.85, 6.56, 11.26 and 16.44 mW. All the other experiments were performed with a light intensity of 11.2 mW.

### Immunohistochemistry

Brain immunofluorescence was carried out as described previously (Chu et al., 2014). Primary antibodies used were chicken anti-GFP (1:1000, Abcam #13970) and mouse anti-brp (1:50, DSHB #nc82). Secondary antibodies used were goat anti-chicken Alexa 488 (1:200, Abcam #150169) and goat anti-mouse Alexa 568 (1:200, Thermo Fisher Scientific #A11004).

All images were acquired using a Leica SP5 II Confocal microscope with a 25x water immersion objective. All images were taken sequentially with a z-stack step size at 2 mm, a line average of 2, speed of 200 Hz, and a resolution of 1024 × 1024 pixels.

### Statistical analysis

Statistical tests were performed using GraphPad Prism 6 software. Descriptions and results of each test are provided in the figure legends. Sample sizes are indicated in the figure legends.

Sample sizes were determined prior to experimentation based on the variance and effect sizes seen in prior experiments of similar types. All experimental conditions were run in parallel and therefore have the same or similar sample sizes. All replicates were biological replicates using different flies. Data for behavioral experiments were performed with flies from at least two independent crosses. There was one condition where data were excluded, which were determined prior to experimentation and applied uniformly throughout: the data from individual flies were removed if the fly did not pass a set minimum threshold of sips (15), or the data showed hallmarks of a technical malfunction (rare).

## Supporting information

## Acknowledgements

We thank Carlos Ribeiro and Pavel Itskov for kindly providing their original FlyPAD VHDL code, which was instrumental in developing the STROBE system. This work was funded by Natural Sciences and Engineering Research Council (NSERC) grants RGPIN-2016-03857 and RGPAS 492846-16. M.D.G. is a CIHR New Investigator and a Michael Smith Foundation for Health Research Scholar.

## Authors contributions

R.C.W.C. and H.Z. designed and built the STROBE with conceptual input from P-Y.M. and M.D.G., and wrote the STROBE methods section. P-Y.M., P.J., M.J. and D.F-K performed behavior experiments under the supervision of P-Y.M. and P.J.. P-Y.M. and P.J. performed immunostainings. P-Y.M and M.D.G wrote the paper with input from the other authors. M.D.G. supervised the project.

## Supplementary materials

**Supplementary Figure 1.**
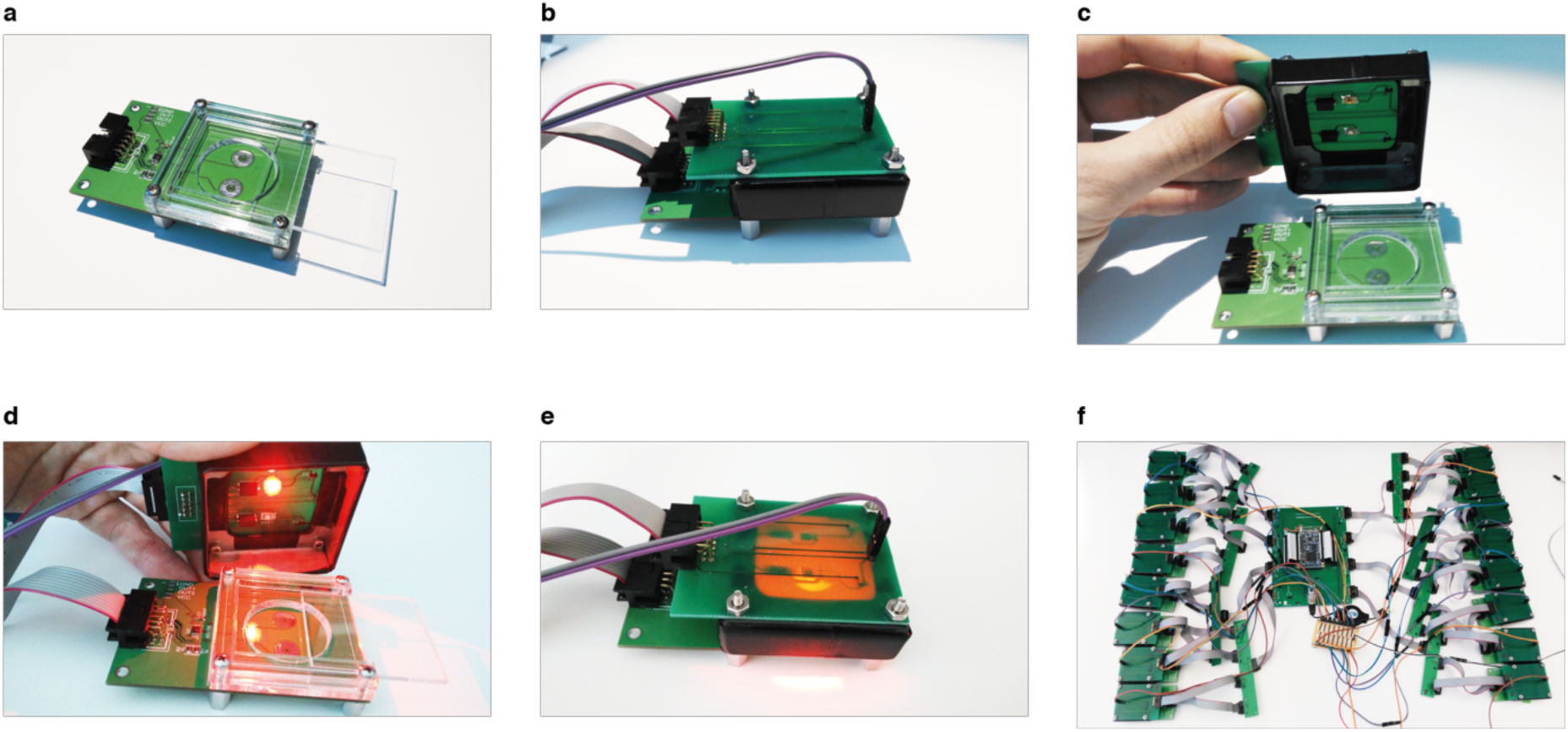
The STROBE setup. **a** FlyPAD arena. **b**-**c** FlyPAD arena with STROBE lid. One LED is positioned just above each channel **d-e** STROBE arena with red LED on. **f** Complete STROBE setup, 16 arenas can work in parallel.

**Supplementary Figure 2.**
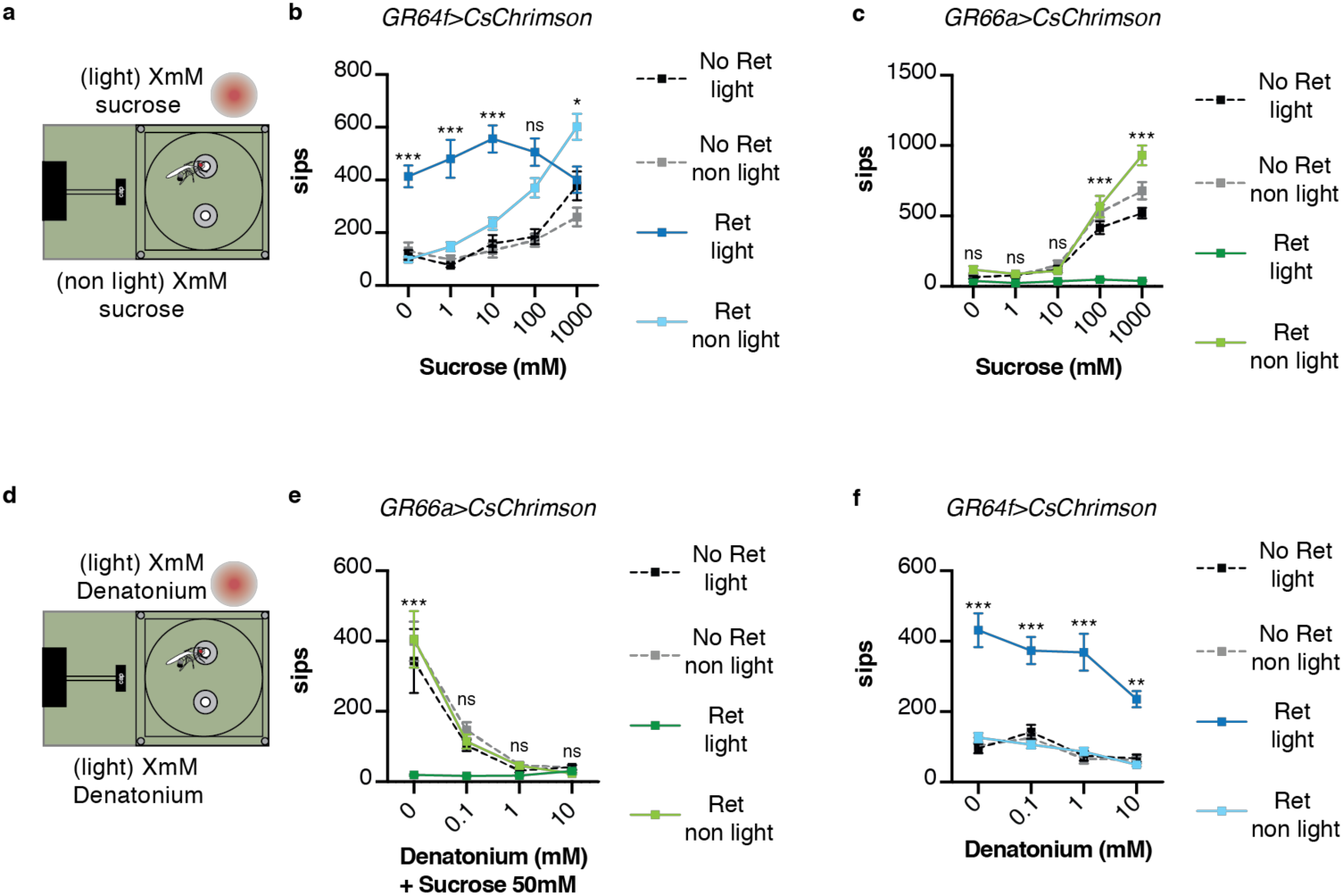
Chemical taste ligands suppress impact of light evoked GRN activity. **a** Experimental setup: both channels contain the same sucrose concentration (0, 1, 10 100, 100 mM) mixed with 1% agar, with one triggering LED activation. **b-c** The effect of sucrose concentration on the number of sips on each channel for *Gr64f>CsChrimson* (b) or *Gr66a>CsChrimson* (c) (b, retinal flies: *n* = 28, 30, 30, 27, 30; control flies: *n* = 25, 26, 22, 25, 21; c, retinal flies: *n* = 28, 34, 34, 26, 46; control flies: *n* = 32, 40, 40, 37, 51). These data are presented as preference indices in Figure 4 b,c. Experimental setup: both channels contain the same denatonium concentration (0, 0.1, 1, 10 mM). For Gr66a>CsChrimson activation, both channels also contain 50 mM sucrose. **e-f** The effect of denatonium concentration on the number of sips on each channel for *Gr66a>CsChrimson* (e) or *Gr64f>CsChrimson* (f) (e, retinal flies: *n* = 25, 27, 30, 31; control flies: *n* = 26, 28, 27, 28; f, retinal flies: *n* = 30, 29, 29, 29; control flies: *n* = 30, 28, 27, 30). These data are presented as preference indices in Figure 4e,f. Means are ± SEM; statistical test: *two-way ANOVA*; *Tukey post-hoc*; ns: *p* > 0.05; *p < 0.05; \*\**p* < 0.01; \*\*\**p* < 0.001.

**Supplementary Figure 3.**
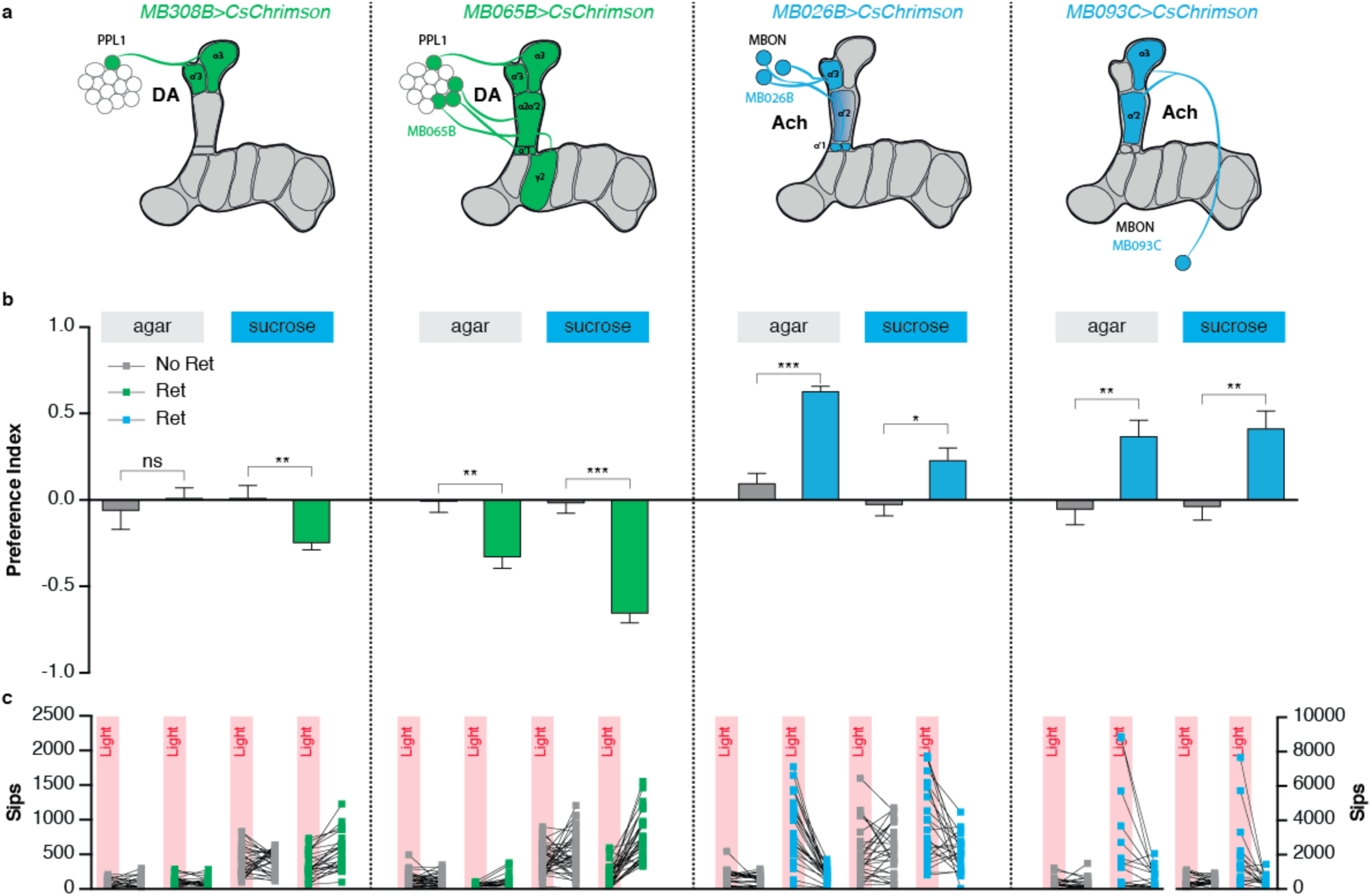
Manipulation of PPL1 DANs and their corresponding output neurons modifies feeding behavior. **a** Schematics of the lines used in the experiment. **b-c** The effect of two PPL1 drivers and two MBON drivers innervating similar compartments on preference (b) and sip numbers (c) in the STROBE. Each experiment is shown with plain agar in both channels, and with sucrose in both channels (MB308B: *n* = 20, 22, 27, 29; MB065B: *n* = 35, 22, 38, 31; MB026B: *n* = 26, 24, 29, 24; MB093C, *n* = 23, 17, 26, 18). Values are means ± SEM; statistical test: *t*-test; ns: *p* > 0.05; \**p* < 0.05; \*\**p* < 0.01; \*\*\**p* < 0.001 in comparison between two groups.

**Supplementary Figure 4.**
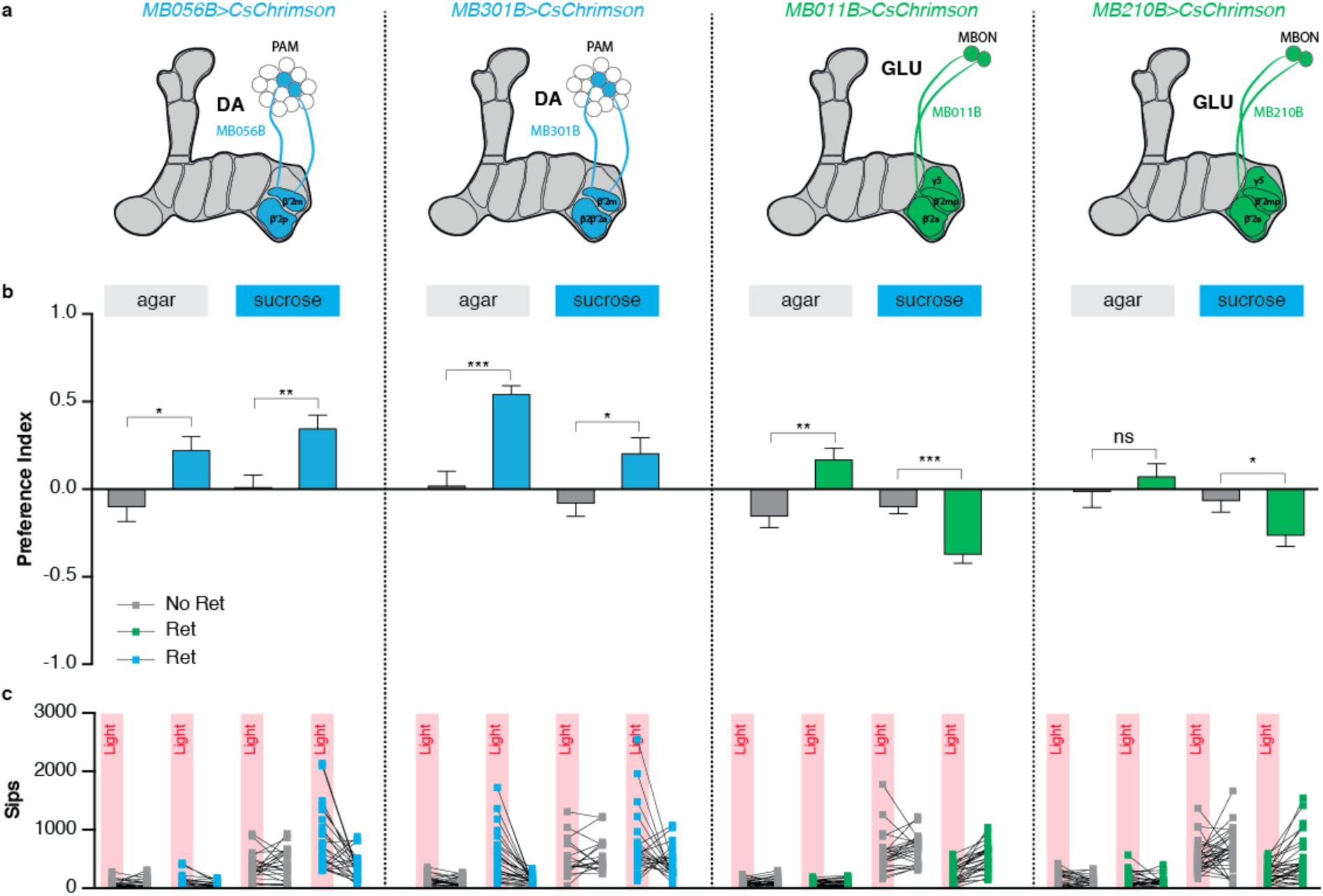
Manipulation of PAM DANs and their corresponding output neurons modifies feeding behavior. **a** Schematics of the lines used in the experiment. **b-c** The effect of two PAM drivers and two MBON drivers innervating similar compartments on preference (b) and sip numbers (c) in the STROBE. Each experiment is shown with plain agar in both channels, and with sucrose in both channels (MB056B: *n* = 15, 16, 24, 24; MB301B: *n* = 25, 28, 18, 24; MB011B: *n* = 29, 27, 25, 27; MB210B, *n* = 24, 28, 30, 31). Values are means ± SEM; statistical test: *t*-test; ns: *p* > 0.05; \**p* < 0.05; \*\**p* < 0.01; \*\*\**p* < 0.001 in comparison between two groups.

**Supplementary Figure 5.**
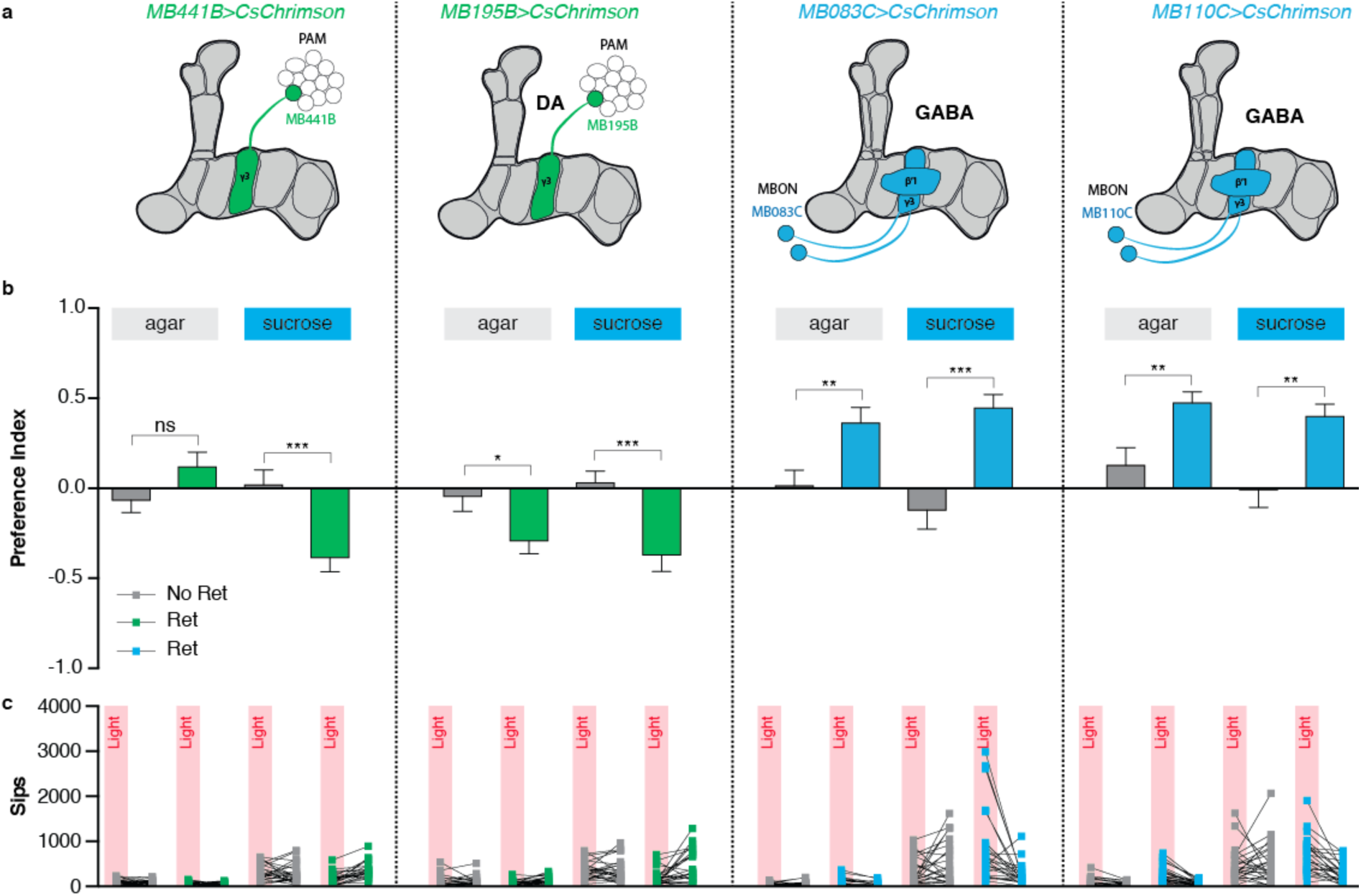
Manipulation of PAM γ3 DANs and their corresponding output neurons modifies feeding behavior. **a** Schematics of the lines used in the experiment. **b-c** The effect of two PAM γ3 drivers and two MBON drivers innervating similar compartments on preference (b) and sip numbers (c) in the STROBE. Each experiment is shown with plain agar in both channels, and with sucrose in both channels (MB441B: *n* = 30, 25, 25, 24; MB195B: *n* = 23, 24, 26, 25; MB083C: *n* = 25, 21, 25, 25; MB110C: *n* = 17, 26, 27, 24). Values are means ± SEM; statistical test: *t*-test; ns: *p* > 0.05; \**p* < 0.05; \*\**p* < 0.01; \*\*\**p* < 0.001 in comparison between two groups.

