## Supplementary material for "The STROBE: a system for closed-looped optogenetic control of freely feeding flies": STROBE Assembly Package Instructions.docx

Assembling the Strobe System

### Foreword

This document is intended to be a set of instructions and directions for fabricating and assembling the STROBE system extension module and accessory components such as the resistor board and power splitter board. This document should be in a folder named “STROBE Assembly Package” along with documents of other technical formats that would be used in the fabrication process.

### Lighting PCB

#### PCB Manufacturing

To manufacture the PCB, specialized facilities are needed. Thus, it is recommended to find a PCB manufacturer and provide the necessary files for them to manufacture the PCBs. In the main folder, there is a subfolder named “Lighting PCB Fabrication”. This folder contains the Gerber (.gbr) files which contain all the information needed for the manufacturer to produce the PCBs.

#### PCB Soldering

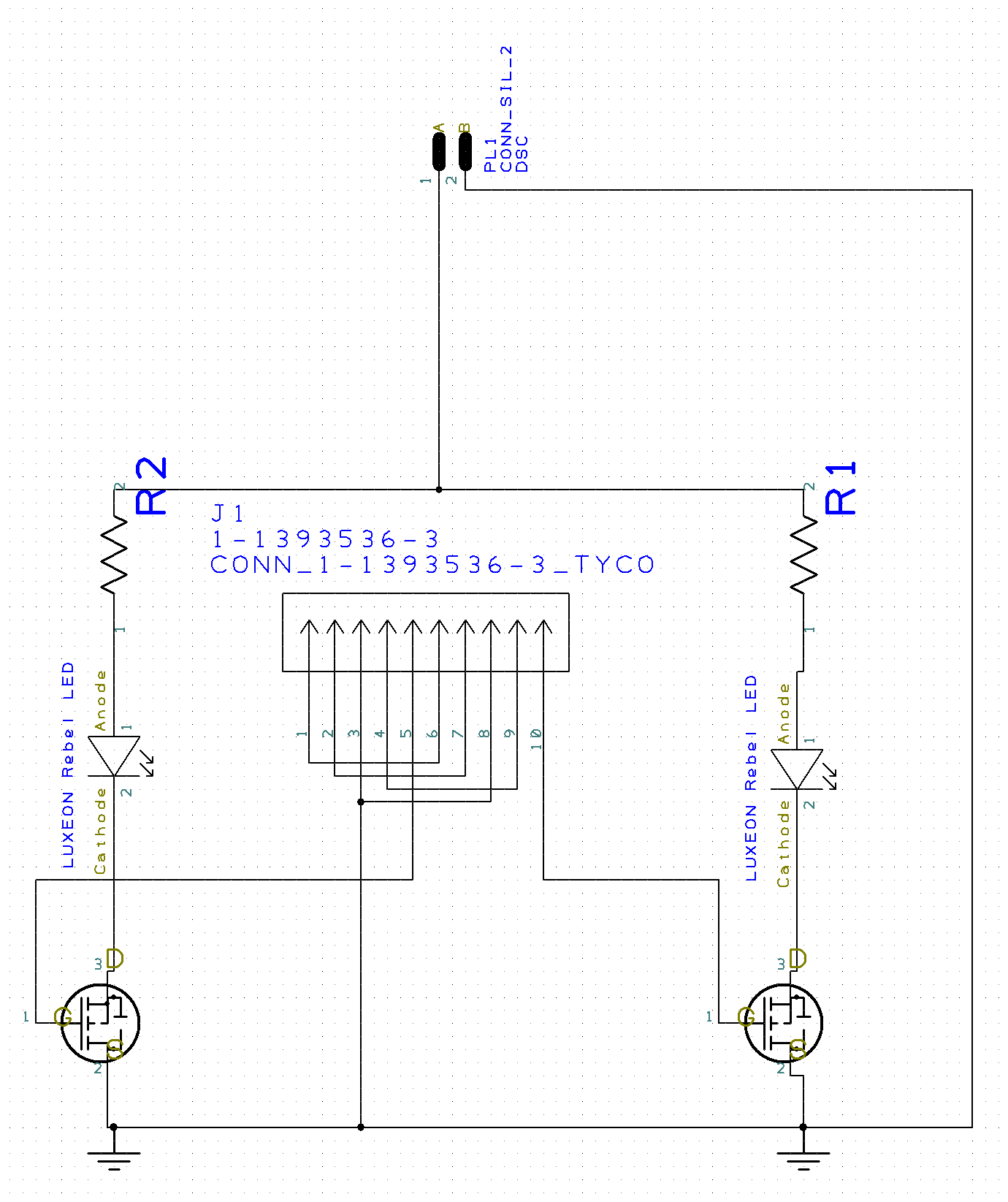

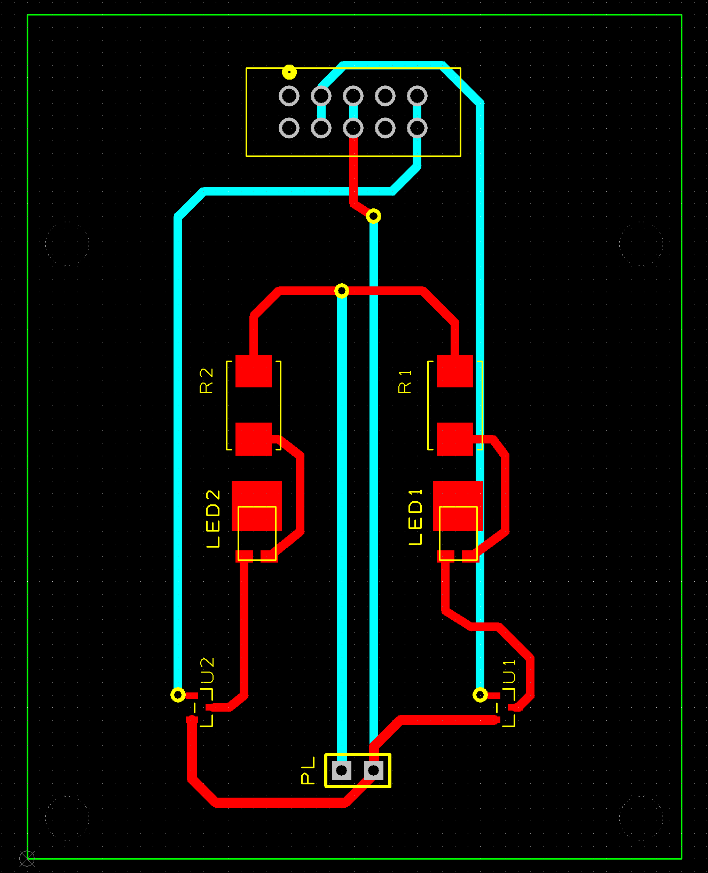

The above two pictures can be found in the folder. The image on the left, named “Lighting Circuit Schematic”, shows the electrical connections of the components on the lighting PCB. The image on the right shows the lighting PCB itself. The blue lines represent copper pathways on the bottom side of the PCB, whereas the red lines represent the pathways on the top side of the PCB. The outlines of the components are in yellow.

The components in the PCB’s design are as follows:

- 2x LUXEON Rebel color LEDs such as (LXM2-PH01-0060 or LXML-PB01-0040)
- 2x Power Resistors (TE Connectivity Passive Product SMW24R7JT)
- 2x N-Channel MOSFETs (Infineon Technologies IRLML0060TRPBF)
- 1x 2-pin header (Like 732-5315-ND on Digikey, these should be abundantly available in any lab with soldering equipment)
- 1x 10-pin connector (3M15452-ND on Digikey)

The person who performs the soldering should be able to tell from the yellow traces on the PCB how the components should be placed and soldered. **Note: the 2-pin and 10-pin header are supposed to go on the reverse side of the PCB.** (All other components go on the conventional side.) The end product should look like this:

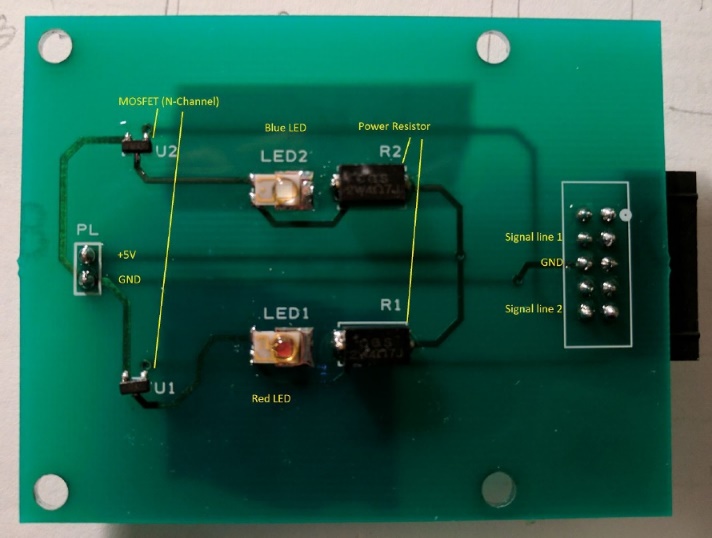

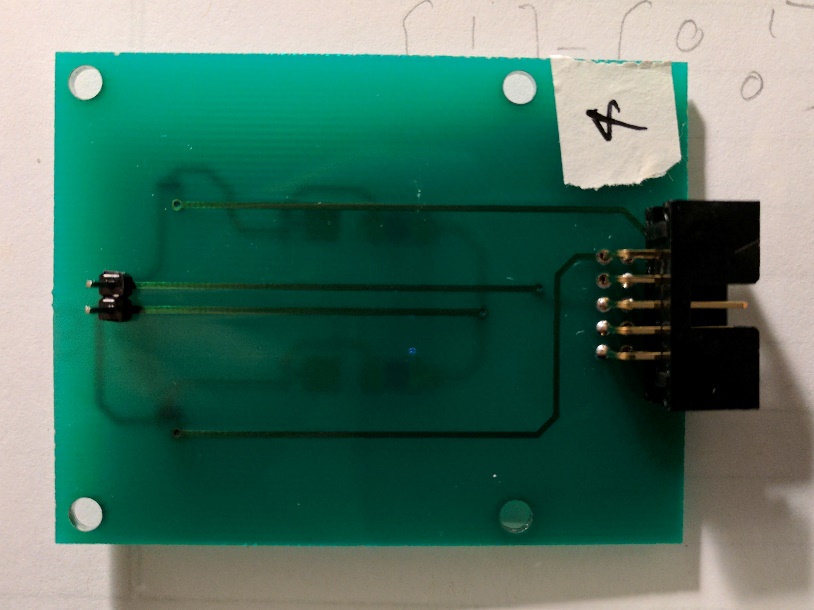

### PCB Housing

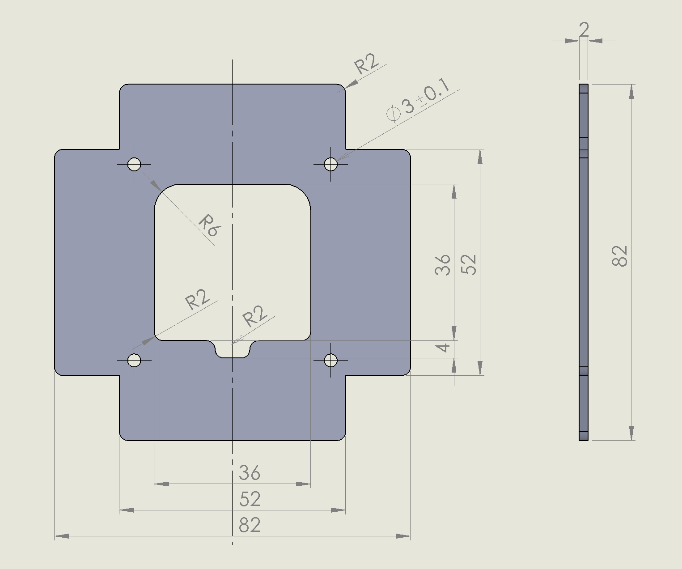

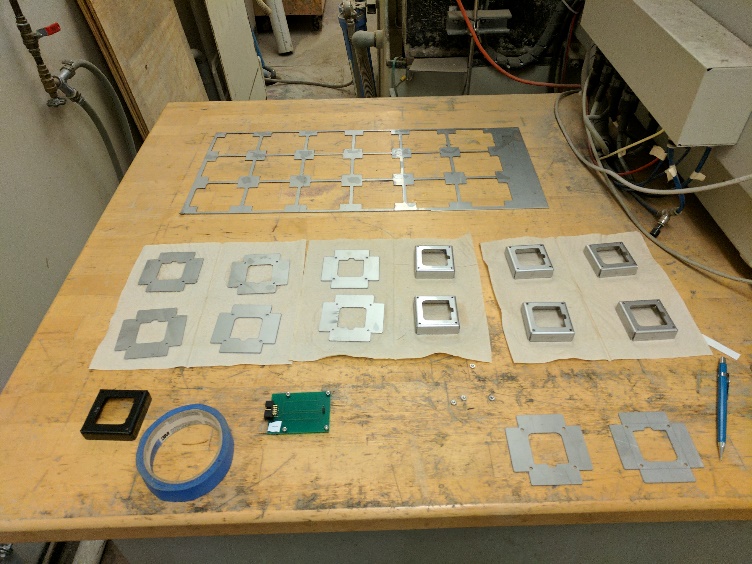

The image on the right is a dimensioned draft of the PCB housing and can be found in the main folder. The image on the right shows how the housing units look in the different steps in the fabrication process with the PCB for scale. The housing part is designed in SolidWorks and the part file (ending in .SLDPRT) is included in the subfolder “Housing Fabrication” along with a flattened pattern .dxf file that can be read by cutting machines like the waterjet.

The PCB housing is manufactured out of 0.05” (1.23mm) stainless steel according to the following steps:

1. Water jetting from steel. The “Flat Pattern Housing.dxf” should be used. The waterjet software should have the ability to add cutting tabs to the pattern, and this should be done to prevent the thin steel from falling into the machine’s crevices mid-cut. The metal sheet should also be weighed down to prevent currents generated from high pressured water to displace the metal sheet.

3. Bend the four tabs with the larger area of the cutout always clamped down. This prevents warping during bending on the most fragile area, the notched metal section. Because of the natural springiness of the steel, the bending press should be brought slightly past 90 degrees so that when the steel springs back a bit, the tab ends up perpendicular to the center section.

4. The bent units should be deburred to remove sharp edges.

5. Gaps in the deburred units can be covered up with black vinyl electrical tape.

The finished housing unit can be attached to the lighting PCB with any screw (imperial or metric size) between the metric sizes M2 and M3.

### Accessory Boards

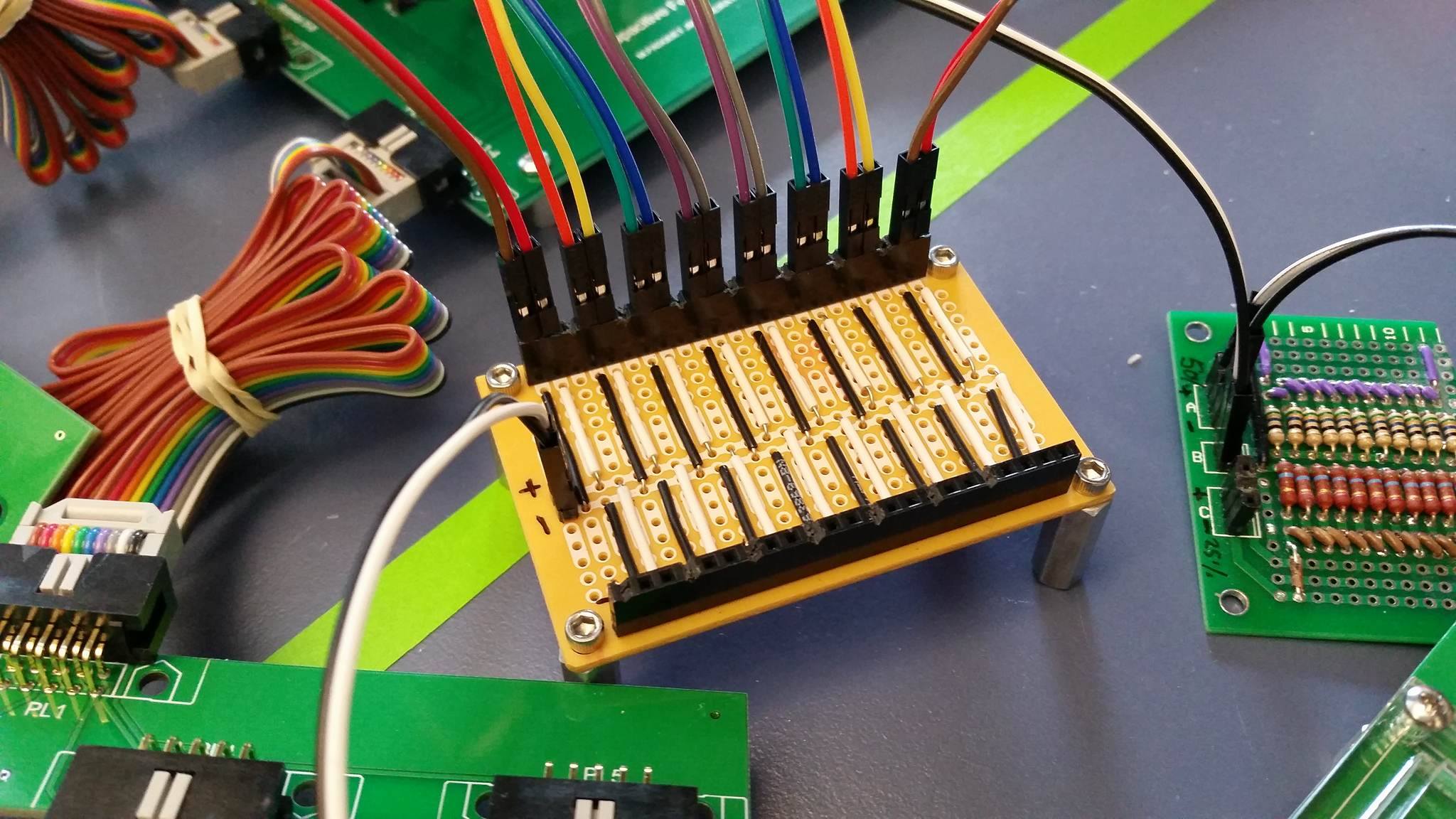

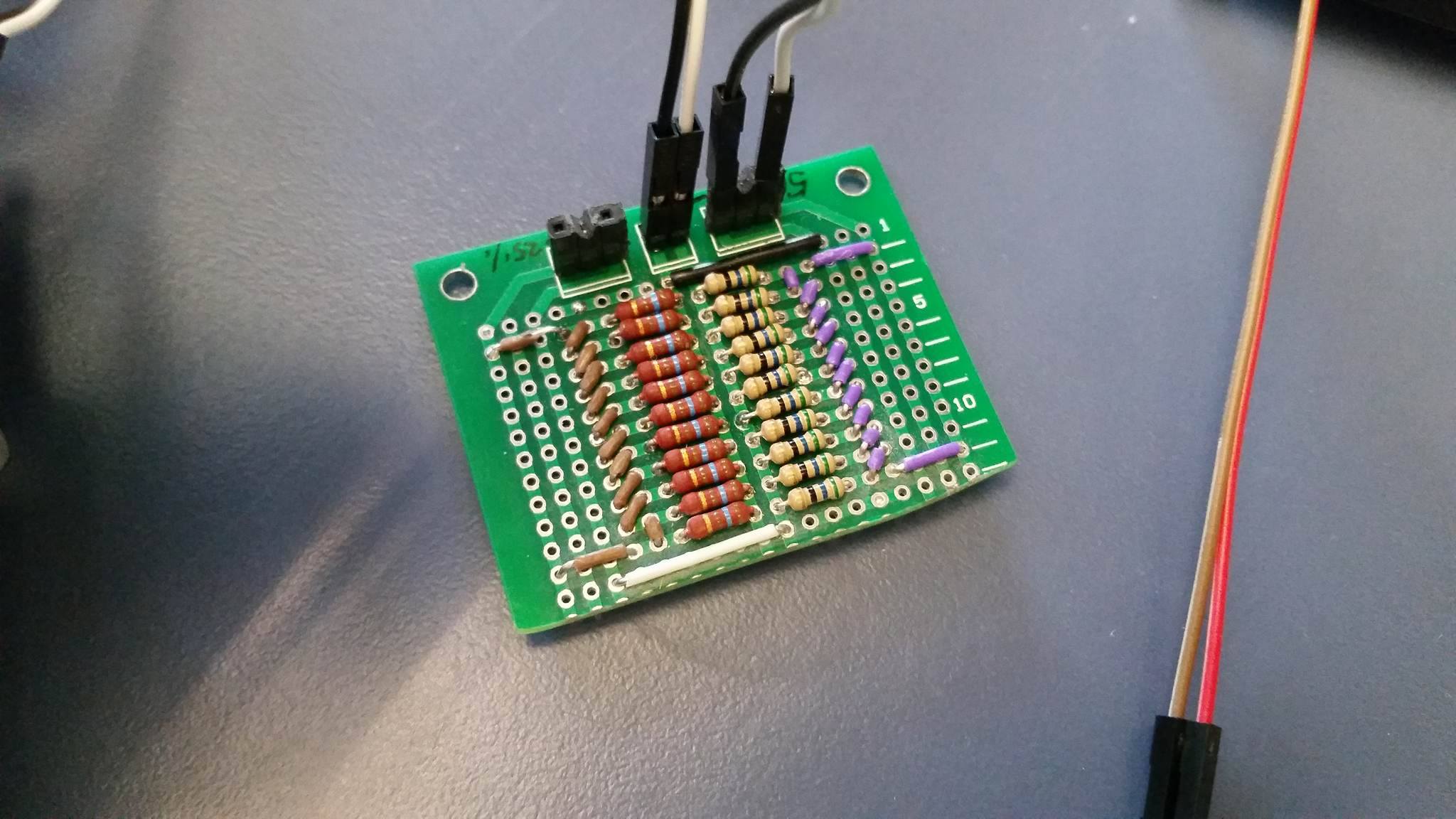

#### Power Splitter Board

The image on the left depicts the power splitter board. This was made on a rather uncommon but perfectly sized prototype board. Any electrical technician or soldering personnel should be able to replicate something with the same functionality if they understand the purpose of this board, since it requires no specific components, just wires and header pins.

The purpose of the board power splitter board is simply to take in a pair of inputs from the STROBE power supply (GND and 5V, labelled – and + on the board in the image) and split this into 16 pairs of outgoing connections, with each pair going to one lighting PCB. One can see the input black and white pair in the picture above being split on the board into 16 pairs of black and white outputs connecting to lighting PCBs via the rainbow pairs of cables.

#### Resistor Board

The image in the above right shows the resistor board, which is used to cut power going to the STROBE system from nominal (100%) to 50% and 25%. Originally this board was made in case some researchers using the STROBE system were not familiar with the power supply settings. However, it does appear that the power supply settings are intuitive, and so this board is not necessary due to the fact that the voltage on the power supply can be adjusted, and gives a continuous range of possible settings.

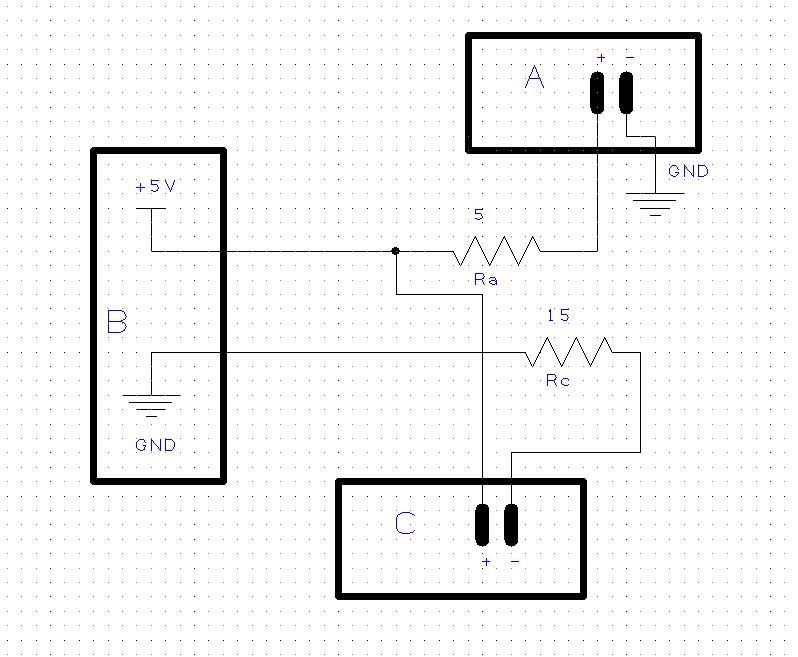

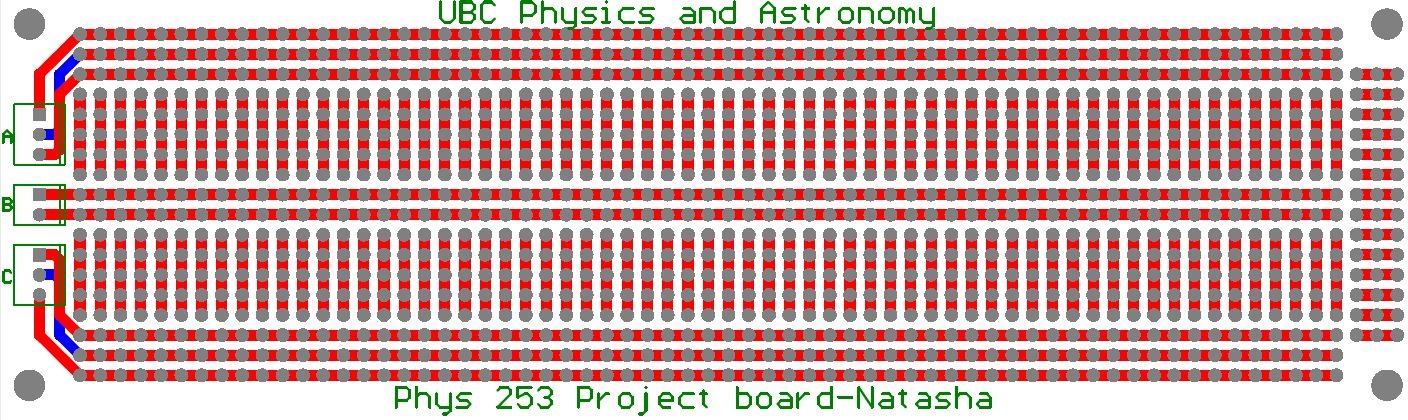

The resistor board circuit design is shown above. This design was originally meant to be implemented on the UBC Physics and Astronomy Department’s Natasha Board (seen in the above right), a prototyping board which had three ports (A, B, C). In this design, the B port takes the power line inputs from the supply. One of ports A or C can be used as an output power line, but not both simultaneously. Depending on which of A or C the output lines are attached to, the LEDs will get 50% or 25% respectively of the current compared to nominal output. In the first resistor board, there are many resistors in parallel which was the only way to handle a larger current load with the parts that were available in the project lab. For future resistor boards however, it is recommended to simply use 4 power resistors like the power resistors on the lighting circuit (5 ohm for the A line and three 5 ohm in series for the C line) on a custom made PCB which also contains the power splitter board. The best solution however is to simply use the power supply to adjust power, eliminating the need for this additional component.

### System Connections

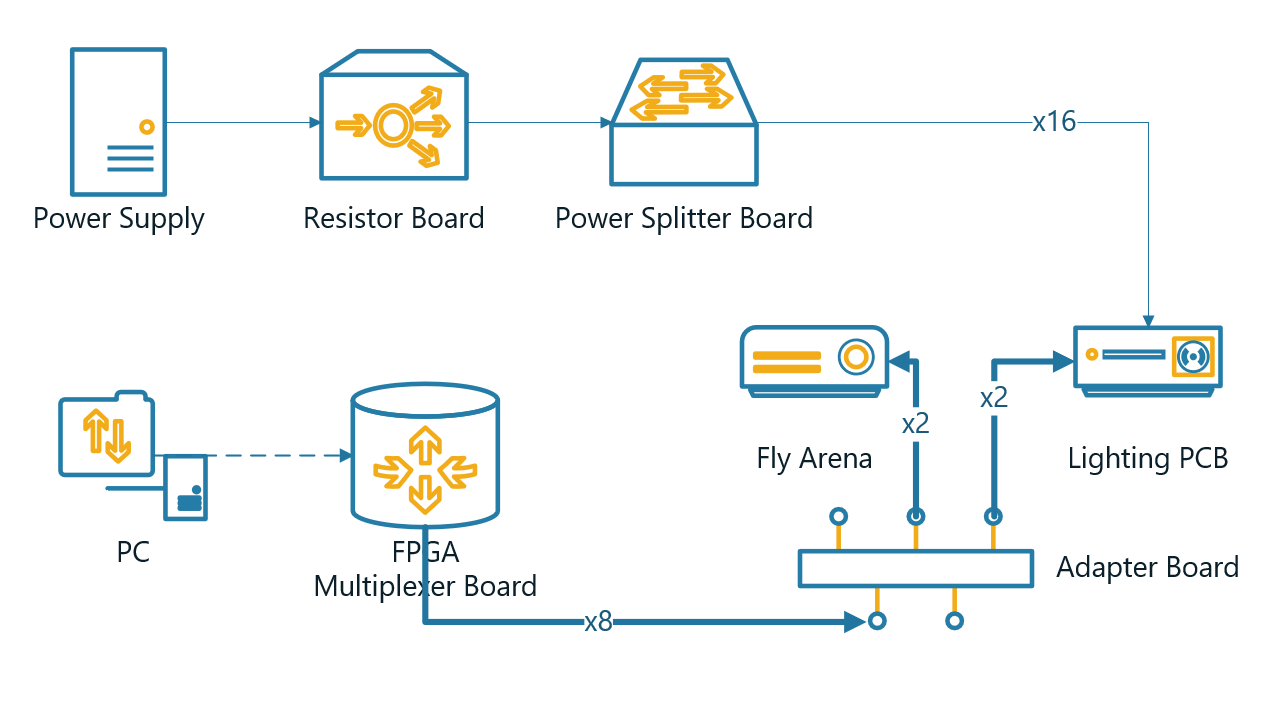

The above diagram shows how the different components of the STROBE system are connected together. The thin arrow connection represents a pair of power lines (+5V and GND), whereas the thick arrow connection represents a 10-pin cable connection. There is also a dashed arrow connection which represents a serial port (USB) connection. Thus, we can see the following connections from the diagram:

1. The power supply connects to the optional resistor board with a single pair of power lines.
2. The optional resistor board connects to the power splitter board with a single pair of power lines. If the resistor board is not used, the power splitter board simply connects straight to the power supply with a pair of power lines.
3. The power splitter board connects to 16 STROBE lighting PCBs, with each connection being a pair of power lines.
4. The PC connects to the FPGA multiplexer board via USB.
5. The FPGA multiplexer board connects to 8 adapter boards each with a 10-pin cable.
6. Each adaptor board connects to two fly arenas and two lighting PCBs each with a 10-pin cable.

Note, with the 10-pin cable, it is important at the adapter board to make sure the fly arenas and lighting PCBs are connected into the right slots. With the power line pairs, it is important to ensure the 5V and GND lines are properly connected and not in reverse.

### Additional Notes

- All the diagrams in this document are included in the STROBE Assembly Package folder.
- Included in the main folder is also the Designspark PCB design files for the lighting PCB. The circuit schematic file is named “Lighting PCB.sch”, and the PCB layout file is named “Lighting PCB.pcb”. The circuit analysis file ending in .asc is also included for future design iteration use.
- The fly arena PCB Gerber files have also been included in the package.
