## Supplementary figures and images for "The STROBE: a system for closed-looped optogenetic control of freely feeding flies"

### Housing Mechanical Dimensions.png

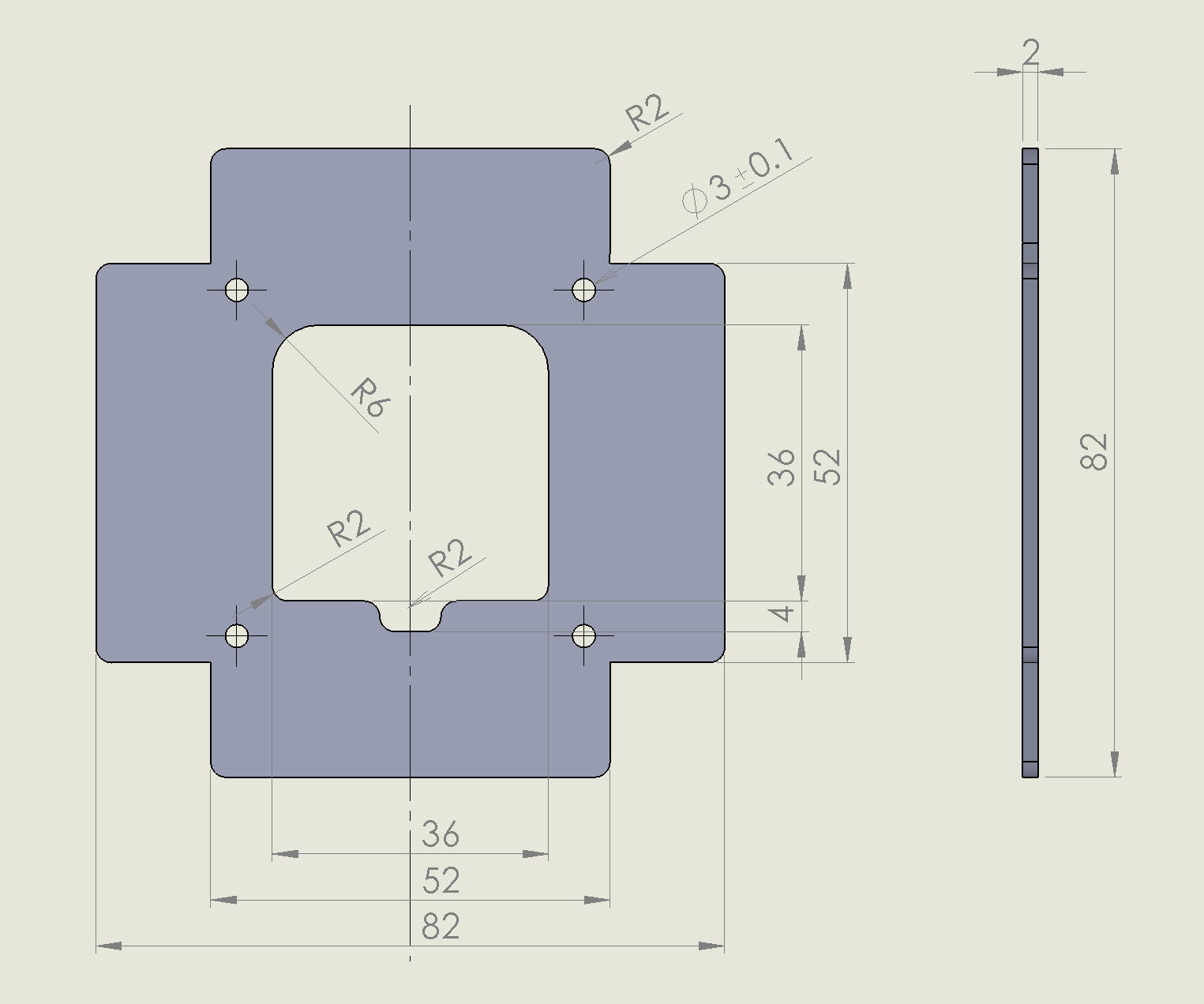

### Lighting Circuit Schematic.png

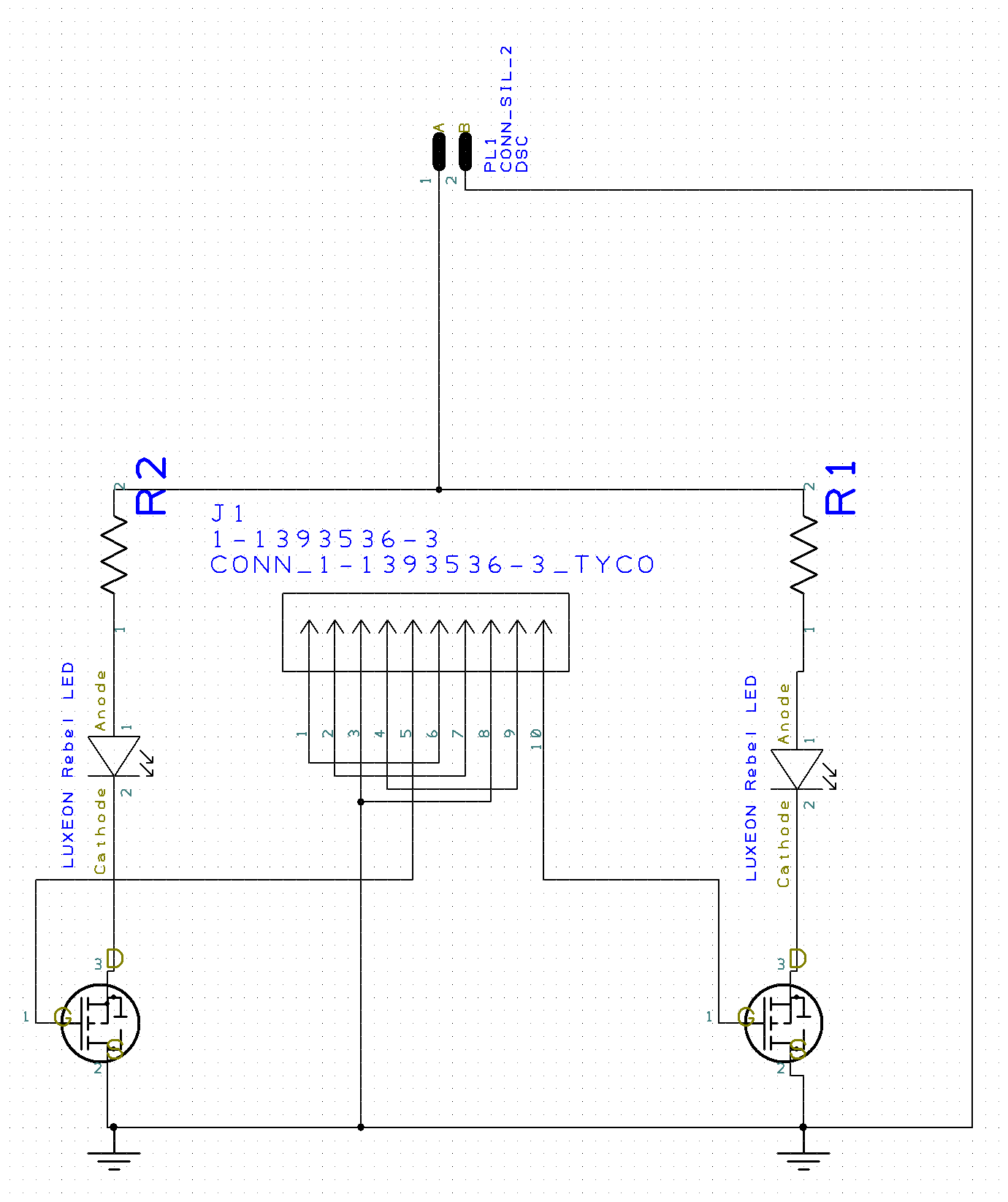

### Lighting PCB Layout.png

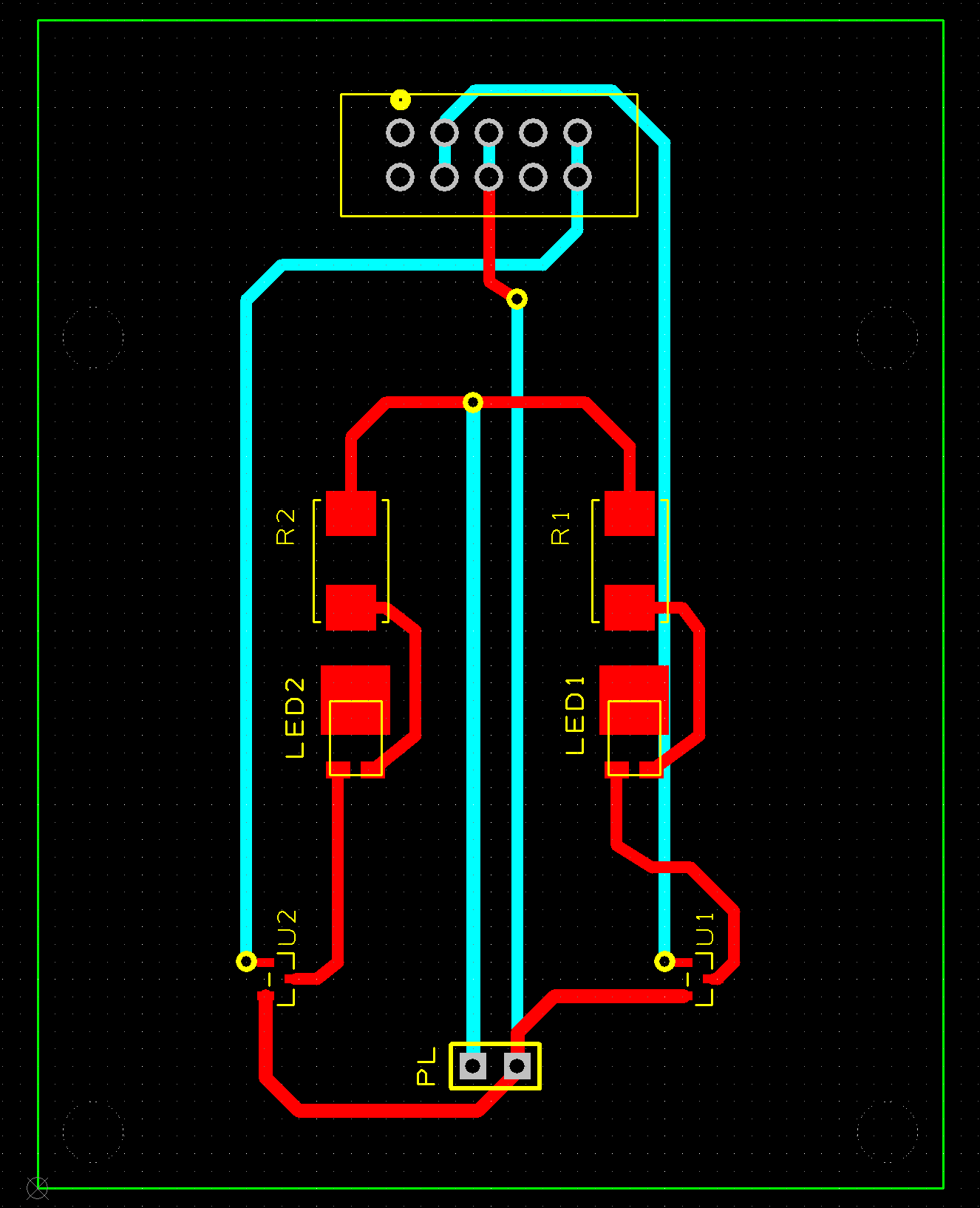

### PCB Housing Fabrication Steps.jpg

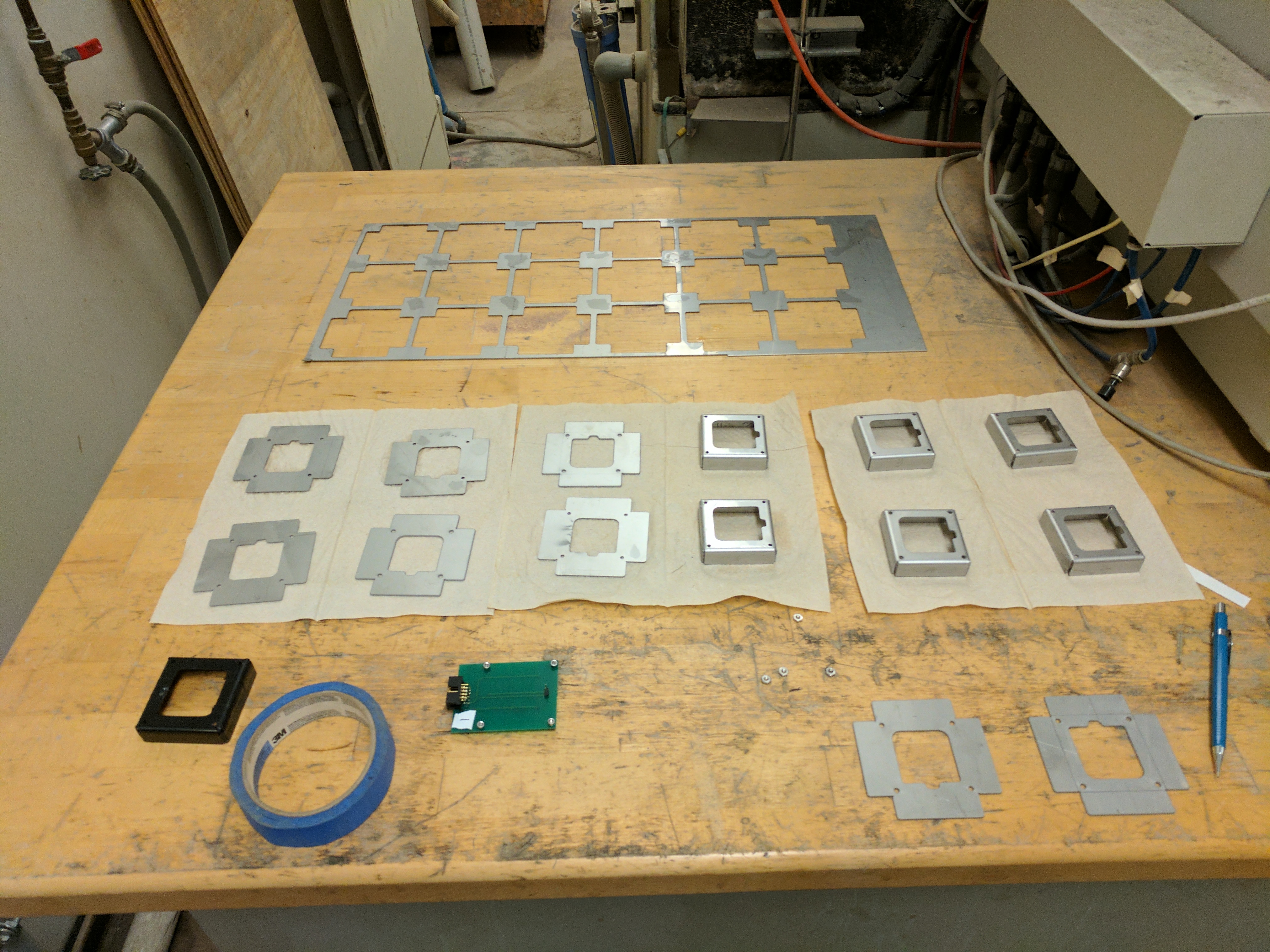

### Power Splitter Board.jpg

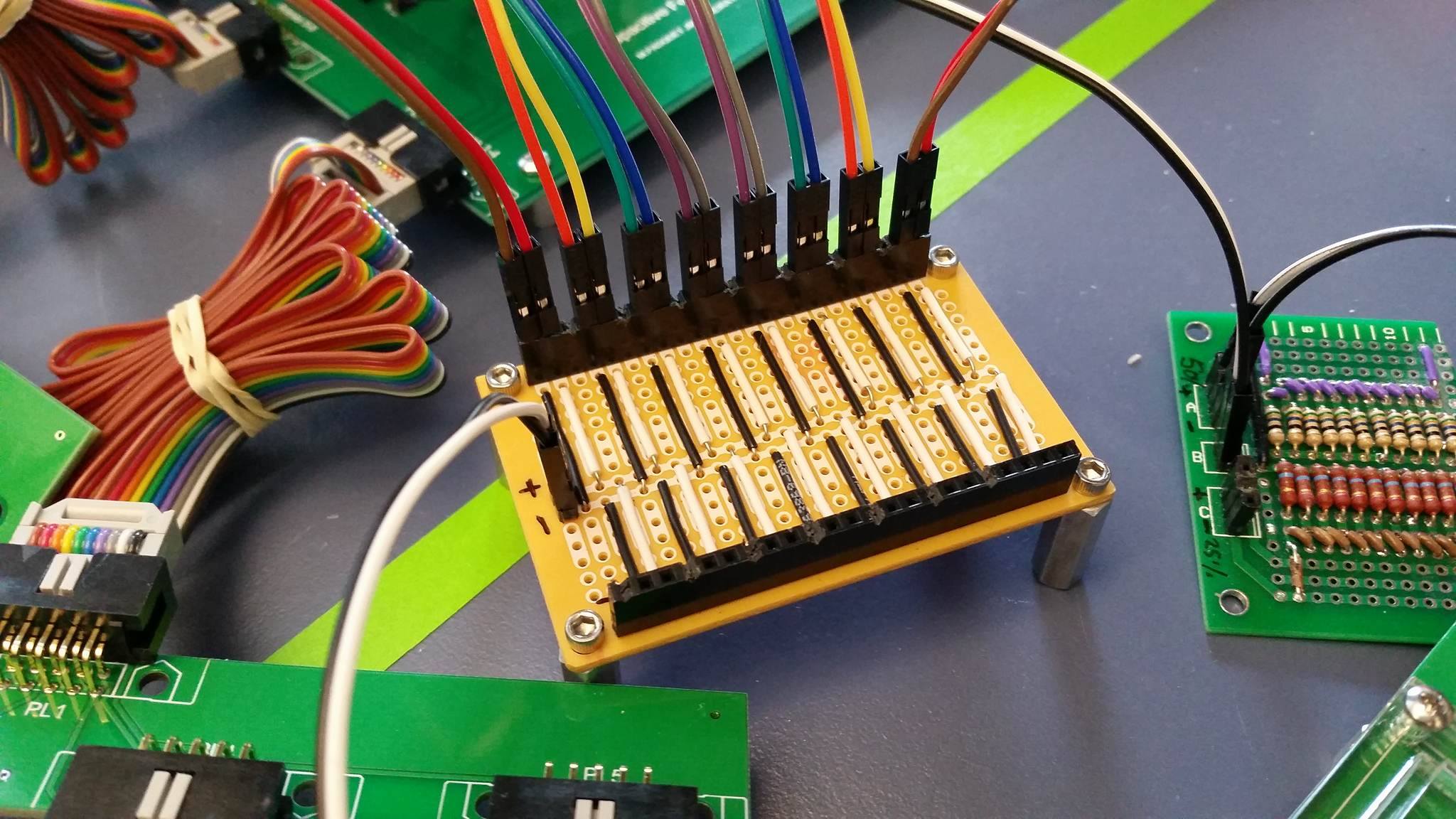

### Project Board Natasha.jpg

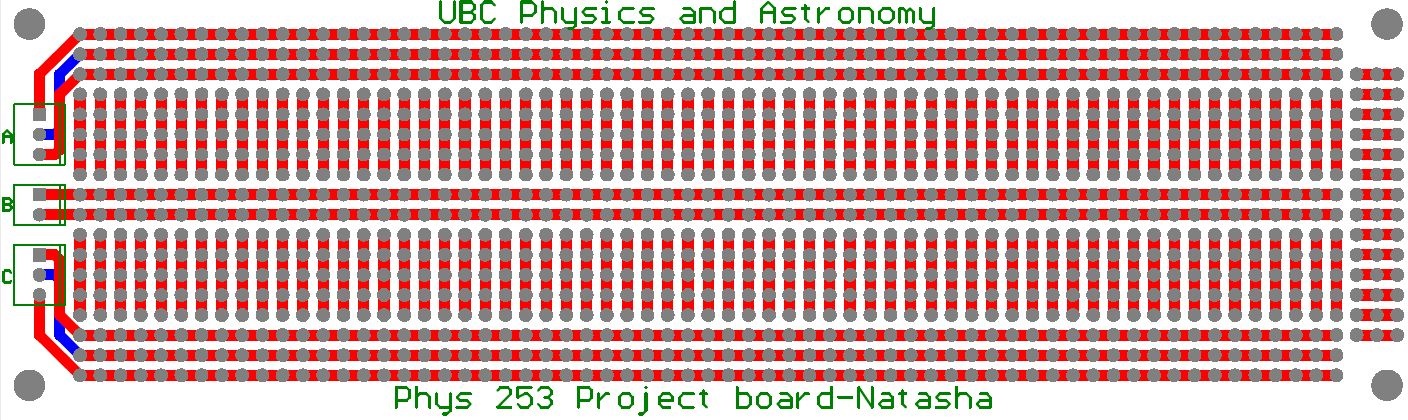

### Resistor Board Schematic.jpg

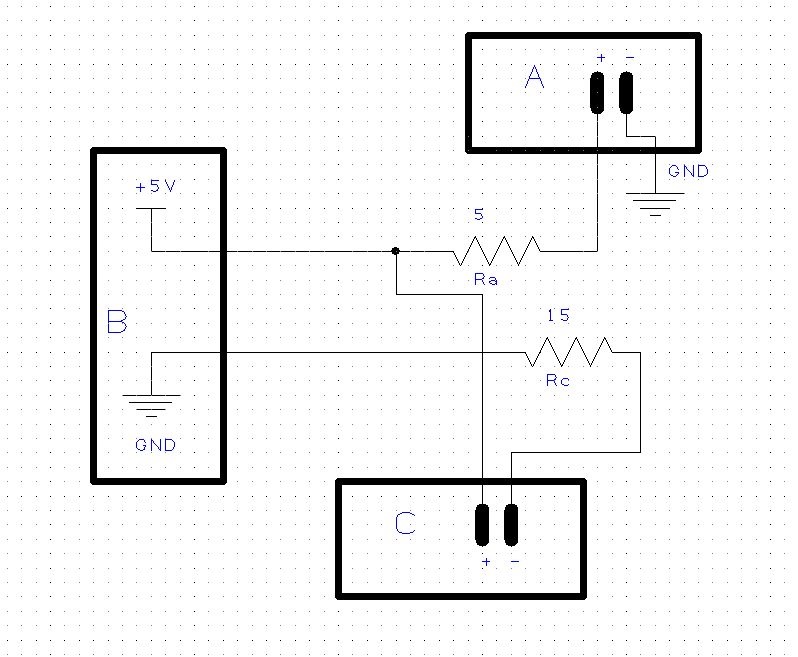

### Resistor Board.jpg

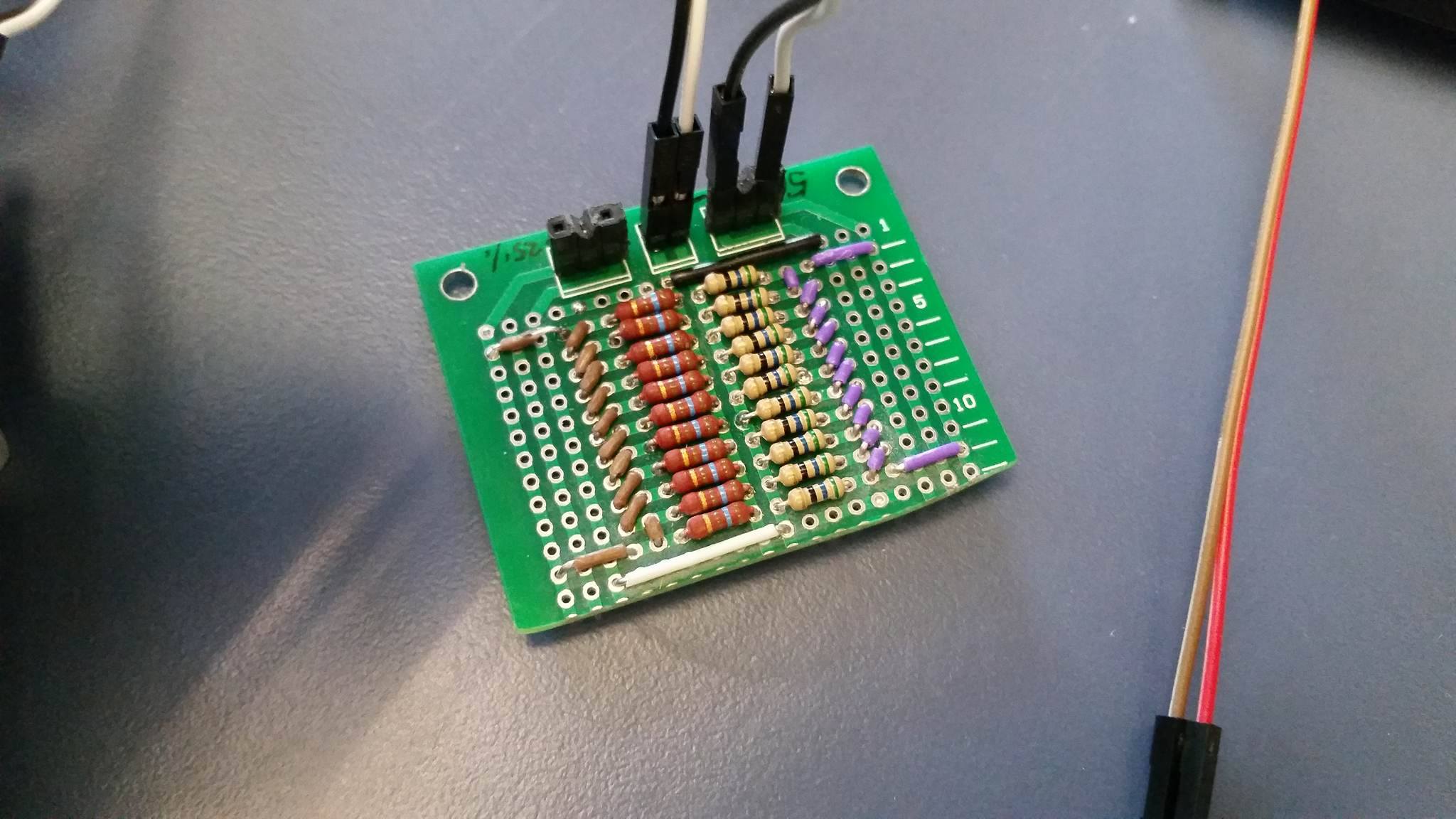

### Soldered PCB Back.jpg

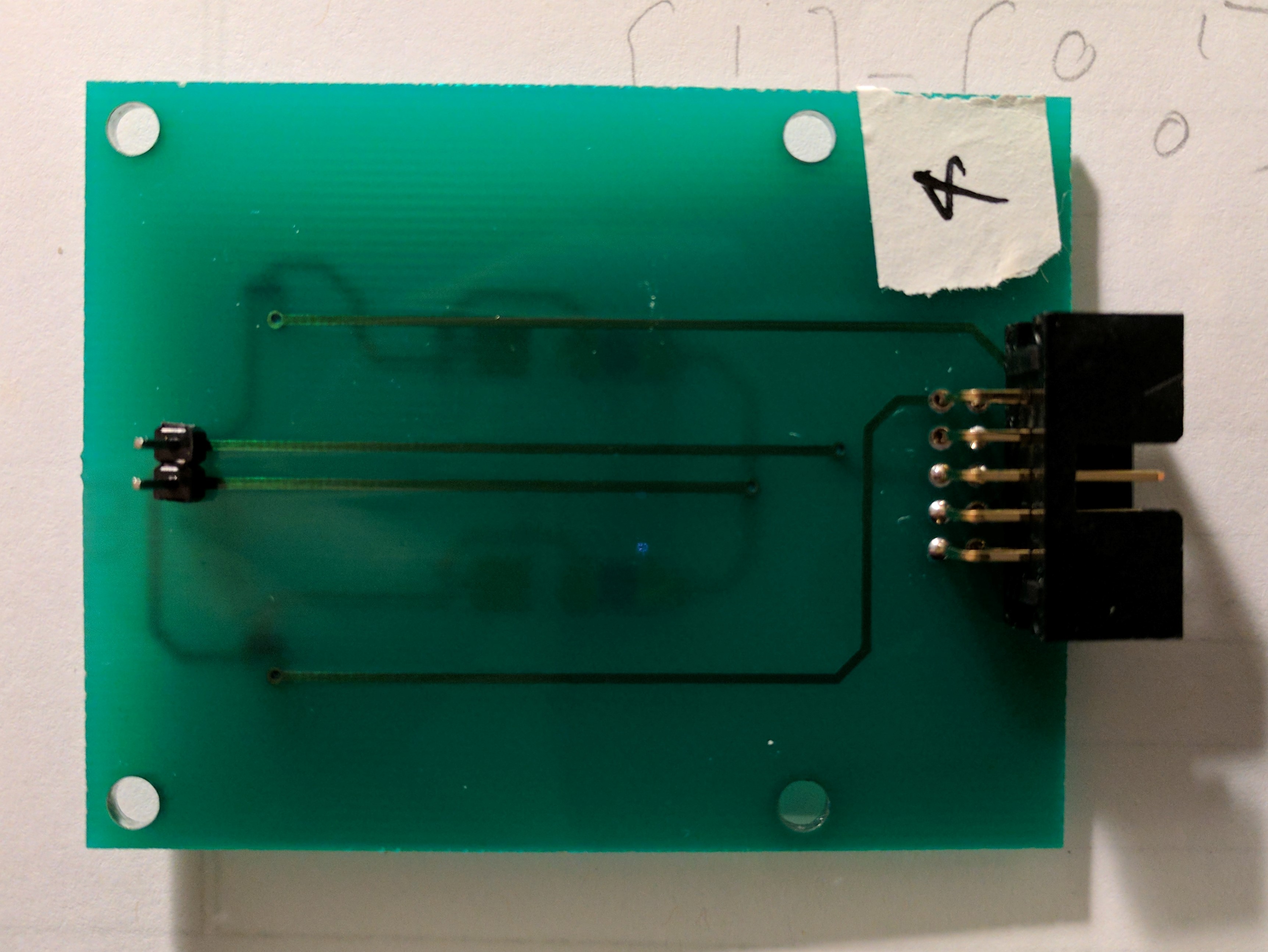

### Soldered PCB Front.jpg

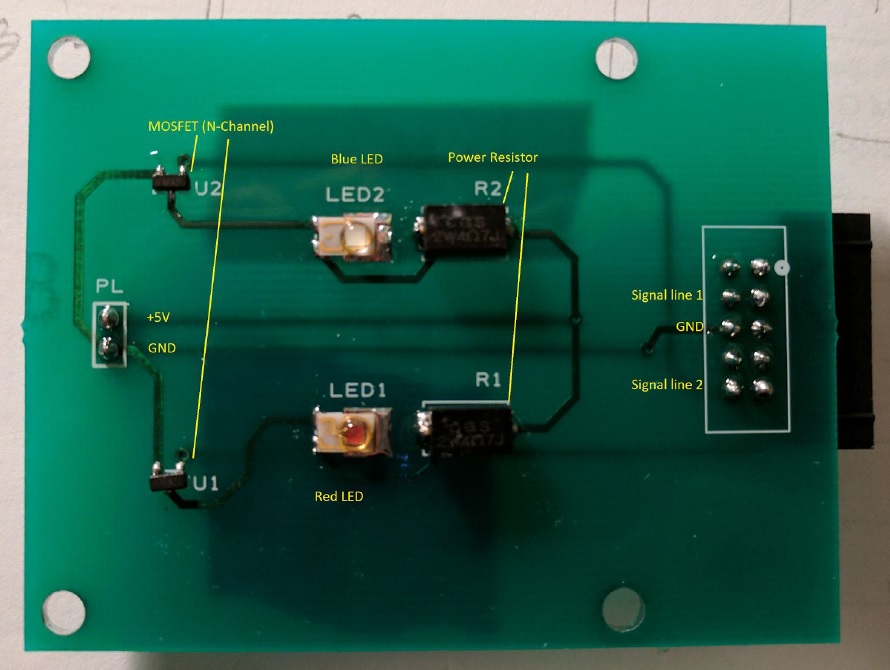

### STROBE System Connections Diagram.png

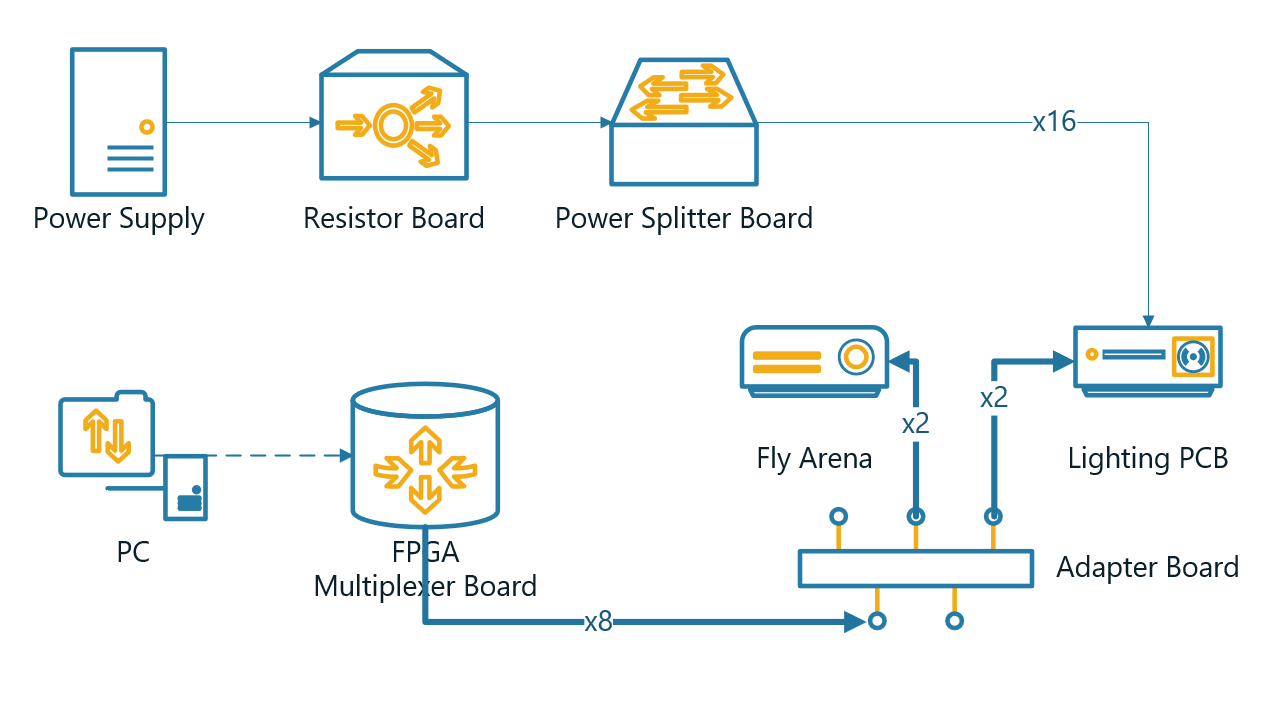
